# A fast-track, high-throughput screening platform for biological nitrification inhibitors discovery based on soil relevant ammonia oxidizing strains

**DOI:** 10.1101/2024.12.03.626636

**Authors:** Alexandros E. Kanellopoulos, Andrea Malits, Hugo Ribeiro, Melina Kerou, Arindam Ghatak, Palak Chaturvedi, Wolfram Weckwerth, Dimitrios G. Karpouzas, Christa Schleper, Evangelia S. Papadopoulou

## Abstract

Nitrogen cycling is critical for ecosystem functioning, with nitrification playing a central role. Excessive nitrification, triggered by heavy nitrogen fertilisation, contributes to reduced nitrogen use efficiency, nitrate leaching, and nitrous oxide emissions. While synthetic nitrification inhibitors (SNIs) help reduce these impacts, their erratic performance and environmental risks have shifted focus to biological nitrification inhibitors (BNIs) as sustainable alternatives. *In vitro* bioassays with ammonia-oxidising microorganisms (AOM) are valuable for BNI discovery and research, but often have low throughput and rely on a limited number of mostly non-soil-relevant or genetically modified ammonia-oxidizing bacteria (AOB) strains, lacking validation with established BNIs. We present a refined fast-track, high-throughput assay for BNI screening, utilizing soil-relevant, ecophysiologically and phylogenetically diverse AOB (*Nitrosospira multiformis, Nitrosomonas ureae, Nitrosomonas communis*) and ammonia-oxidizing archaea (AOA) strains (*Nitrososphaera viennensis,* “*Ca*. Nitrosocosmicus franklandianus”), achieving consistent cellular activity and density. The assay was validated with established SNIs and BNIs, showing differences in inhibition efficacy and strain sensitivity, consistent with literature. As a proof of concept, root exudates from diverse wheat genotypes were screened, demonstrating distinct inhibition profiles. The proposed system advances previously available screening systems and, when integrated with realistic soil tests, will facilitate the discovery of novel BNIs and BNI-producing plant genotypes.

## Introduction

The nitrogen cycle consists of various microbially-mediated processes, with nitrification being the rate-limiting step (1). Ammonia oxidation, catalysed in soil by ammonia-oxidizing microorganisms (AOM), such as ammonia-oxidising bacteria (AOB) from the genera *Nitrosospira* and *Nitrosomonas*, and ammonia-oxidizing archaea (AOA) from the genera *Nitrososphaera, Nitrosotalea* and *Nitrosocosmicus* (2–4), sets the pace for nitrification (3,4). Additionally, “Comammox” *Nitrospira* bacteria, capable of oxidizing both ammonia and nitrite, have been recently discovered (5,6).

The nitrogen cycle plays a key role in converting inert dinitrogen gas into reactive forms essential for biomass production. However, excessive use of the Haber-Bosch process (7,8) has dramatically increased anthropogenic soil ammonia input (7,9), disrupting natural nitrogen fluxes and exacerbating nitrification and denitrification, with detrimental environmental and agricultural implications. Nitrification contributes to low nitrogen use efficiency (NUE), as ammonia (NH_3_) or ammonium (NH_4_^+^) from fertilisers is oxidized to nitrate (NO_3_^-^) (10,11), a molecule prone to leaching from agricultural soils (12). This contributes to eutrophication of aquatic ecosystems (11) and contamination of drinking water resources (13). Furthermore, nitrification and incomplete denitrification are primarily responsible for nitrous oxide (N_2_O) emissions (10,14,15), while nitrification intermediates such as hydroxylamine and nitrite, can also generate N_2_O (14,16,17), depleting NH_4_^+^/NO_3_^-^ nitrogen.

Nitrification inhibitors (NIs), chemicals that decelerate nitrification (18–20), have significant potential for mitigating the above issues. Synthetic NIs (SNIs), such as 2-chloro-6-(trichloromethyl) pyridine (nitrapyrin), dicyandiamide (DCD) and 3,4-dimethylpyrazole (phosphate) (DMPP), are routinely used in agriculture (21), and recent studies have identified novel SNIs like ethoxyquin (22), substituted triazoles (23), azoles and diazoles (24). However, SNIs have been shown to adversely affect non-target soil microorganisms, including those indirectly involved in nitrification and other, unrelated, microbial populations (25,26). Coupled with their high synthesis cost, environmental pollution risks, and potential to enter the food chain (27), alternative approaches need to be developed.

Biological nitrification inhibition (BNI) relies on the ability of certain plants, under low NH_4_^+^ levels, to exude compounds that inhibit nitrification (27,28). These compounds, known as biological nitrification inhibitors (BNIs), are considered an environmentally friendly alternative to SNIs (11,25,28). Examples include sakuranetin (29) and methyl 3-(4-hydroxyphenyl) propionate (MHPP) (30) from sorghum (*Sorghum bicolor*) roots, and 1,9-decanediol (31) from rice (*Oryza sativa*). BNIs can inhibit nitrification *in vitro* (32,33), while BNI-producing plants can reduce NO_3_^-^ production and greenhouse gas (GHG) emissions in agricultural settings (34,35). BNI incorporation in agricultural practice via plant- or chemical-mediated strategies represents a promising approach, and rigorous screening of candidate BNI-producing plant genotypes, and candidate BNI compounds, will unravel the full potential of BNI and enhance its implementation in modern agriculture.

To assess NI potency, culture-dependent approaches, testing the inhibitory potential of pure compounds or plant exudates on axenic AOM liquid cultures have been employed. These tests enable the calculation of inhibitory endpoints for ammonia oxidation, like the effective concentration 50% (EC_50_) (33,36,37) and the inhibition concentration 80% (IC_80_) (31,32), providing inhibitory thresholds for several potential BNIs (32,33). However, their low-throughput, due to the low specific growth rates of AOM, and high space, biomass and time requirements, has prompted the development of fast-track screening systems. These systems facilitate the rapid identification of NI activity in plant rhizodeposits (38,39), pure plant-derived (29,31,34), and synthetic compounds (24). However, these approaches, relied on a genetically modified strain (38,40) or wild-type strains (31) of a single AOB species of little ecological importance in soil ecosystems, *Nitrosomonas europaea* (32,33). More soil-relevant AOB strains (41) and *Nitrososphaera viennensis,* as a sole soil AOA representative (24), have since been used in NI testing. While these inclusions are an upgrade, the need for testing against ecologically relevant AOA strains has yet to be addressed, compromising the robust assessment of BNI potency, as AOA exhibit varying sensitivities to NIs (32,33,37), distinct ecophysiological diversity and, in many cases, outnumber (3,12,42) and even outperform (12) AOB in soil.

Overall, the low throughput of liquid batch cultures and limited representation of ecologically relevant AOB and AOA strains, in current high-throughput systems, highlight the need for a standardized, ecologically valid, fast-track system to accelerate BNIs discovery. We developed a rapid, miniaturised assay, with a turnaround time of less than a day, using a simple 96-well plate format, for high-throughput screening of candidate BNIs, pure molecules and plant extracts. We included five diverse, soil-derived AOB and AOA strains, generated concentrated AOM cultures, and standardized their behaviour by monitoring cell activity and density. The accuracy of the system was validated by generating EC_50_ values for known SNIs and BNIs and comparing them with inhibitory thresholds from literature, using liquid batch culture approaches as benchmarks. Finally, as proof of concept, we applied our system to screen root exudates from geographically diverse wheat genotypes for BNI activity.

## Materials and Methods

### AOM strains and growth conditions

*Nitrosospira multiformis* ATCC25196 and *Nitrosomonas ureae* Nm10 (provided by Graeme Nicol École Centrale de Lyon, France), were grown in Skinner and Walker (SW) medium (43). *Nitrosomonas communis* Nm2 (obtained from Lisa Stein (University of Alberta, Canada) was grown in a medium according to Koops et al. (44) (from here on NCOM medium), adapted to a 3mM NH_4_^+^ substrate and with phenol red instead of cresol red as pH indicator. All AOB cultures were grown statically, in the dark, at 28°C, in 30 mL Sterilin™ sterile polystyrene bottles (ThermoFischer Scientific, Waltham, Massachusetts, USA) at a working volume of 20 mL.

Axenic liquid cultures of the terrestrial AOA *Nitrososphaera viennensis* EN76 and “*Candidatus* Nitrosocosmicus franklandianus C13” were grown in freshwater medium (FWM) buffered with HEPES to pH of 7.5 and supplied with 2mM NH_4_^+^ (NH_4_Cl) following Reyes et al. (45) with slight modifications. *N. viennensis* was incubated without vitamin solution, but with 1 mM pyruvate to scavenge reactive oxygen species. “*Ca.* N. franklandianus” was cultured with 1mM thiamine HCl and without pyruvate. The AOA were grown in the dark in sterile Duran® borosilicate bottles or 30 mL polystyrene tubes, filled up to two-thirds and shaken at 80 rpm.

### Nitrification Inhibitors

The following NIs were used to assess the performance of the fast – track system. For SNIs, the hydrophilic 3,4-dimethylpyrazole phosphate (DMPP) (46) and the hydrophobic nitrapyrin (47) and 1,2-dihydro-6-ethoxy-2,2,4-trimethylquinoline (ethoxyquin) (37) were included. Regarding BNIs, the hydrophilic methyl 3-(4-hydroxyphenyl) propionate (MHPP) (30) and the hydrophobic sakuranetin (29) and 1,9-decanediol (31) were tested. Providers and purities of analytical standards are summarised in Table S1.

### Development of the fast-track testing system

The workflow of the AOB and AOA fast – track system is presented in Figure 1. Regarding AOB, three 60-mL fractions of cultures at the late logarithmic phase were filtered through hydrophilic 0.22 μm polyethersulphone (PES) membrane filters (Frisenette ApS, Knebel, Denmark), the filter was rotated upside-down and bacterial cells were detached through tapping of the filter for 60 sec. Subsequently, 2 mL of sterile SW medium for *N. multiformis* and *N. ureae*, or sterile NCOM medium for *N. communis*, were passed through the filter to resuspend the remaining cells. The filtrates were centrifugated at 11 000 g for 10 min at room temperature. 1.8 mL of the supernatant was discarded, and all inocula were combined into a final volume of 2 mL. The concentrated inoculum was then diluted with sterile SW or NCOM medium, at ratios of 1:10 for *N. multiformis* and *N. communis* and 0.7:10 for *N. ureae*, gently mixed and 200 μL of the culture were dispersed to the wells of a cellGrade^TM^ 96 – well plate (BRAND GMBH & CO KG, Wertheim, Germany). The plate was incubated with a lid, under the same conditions as the respective AOB cultures. Nitrite production was determined colourimetrically at 540 nm by diazotizing and coupling with Griess reagent (48) at specific time-points for 24 hours, or until sufficient consumption of the ammonium substrate in the control treatments (0.1% v/v ddH_2_O or DMSO) was observed. Samples for measuring the abundance of the *amoA* gene were collected at specific time-points as outlined in Figure 1.

**Figure 1.**
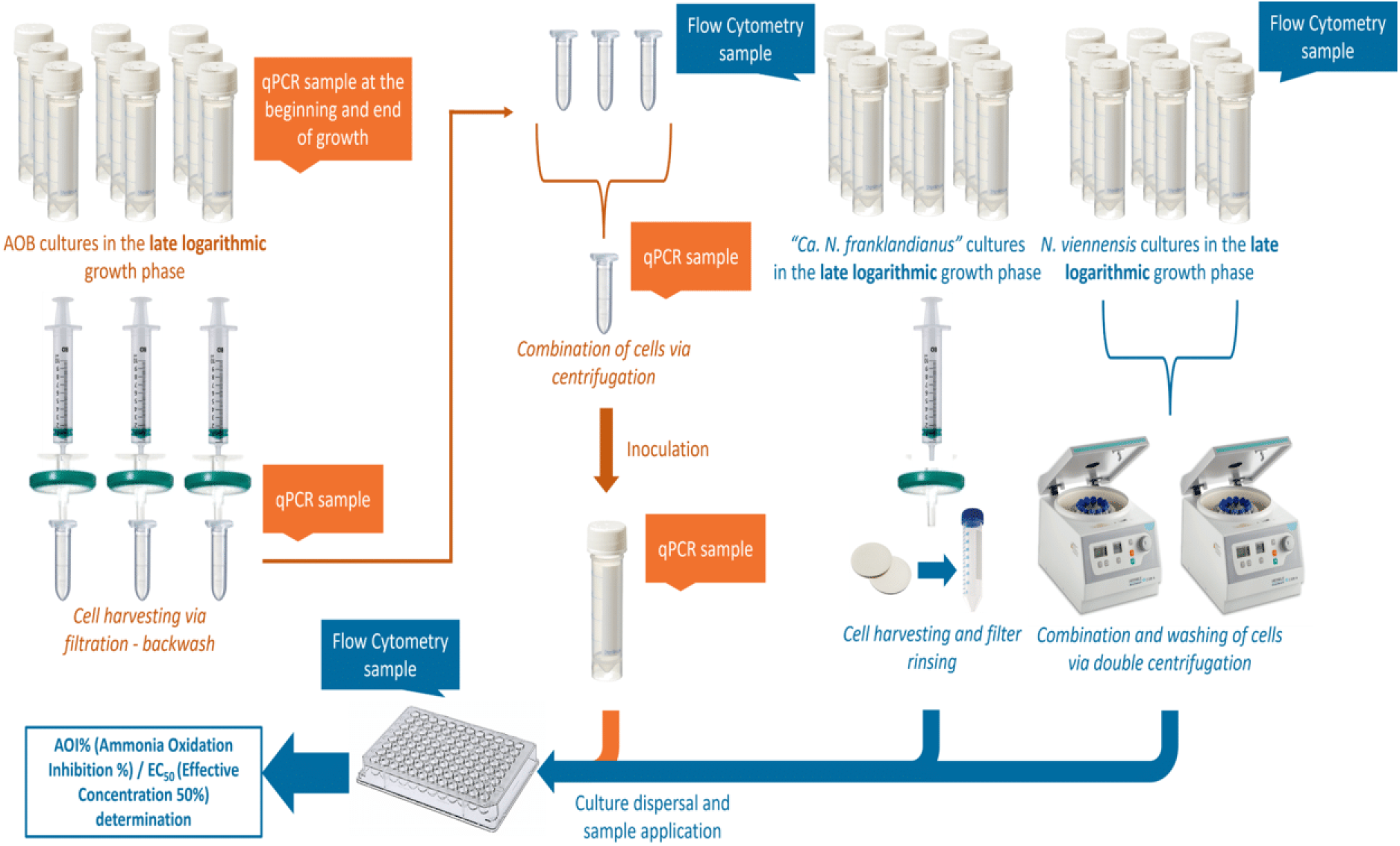
Overview of the development of the fast track, high-throughput system. The AOB workflow is presented in orange, while the AOA workflow is shown in blue. The points of DNA sample collection for qPCR during the procedure specifically refer to the standardization step.

For AOA, *N. viennensis* cultures at the late logarithmic phase were centrifuged at 21 000 g for 30 min at 4°C. The supernatant was discarded, and the pellets resuspended with FWM and centrifuged again to wash out residual nitrite. Again, the supernatant was discarded, and the pellets were resuspended in FWM and kept at 4°C until further processing. “*Ca.* N. franklandianus*”* cultures were concentrated on Millipore® sterile hydrophilic 0.22 μm mixed cellulose esters (MCE) membrane filters using a sterile filtration unit. The cells on the filter were rinsed with FWM to wash the residual nitrite, the filter was placed into a sterile centrifuge tube and FWM was added to achieve the desired cell density. The tube was shaken to detach the cells from the filter, the filter was removed, and the culture concentrate was stored at 4°C until further processing. AOA culture concentrates were allowed to prewarm at 42°C before 200 µL were added in the wells of a 96-well plate, incubated with a lid, statically, at 42°C. Samples were taken at the beginning of incubation and at several time points, until approximately half of the provided NH_4_^+^ substrate was consumed in the control treatments (0.1% v/v ultra-pure water or 0.02% v/v DMSO), to quantify NO_2_^-^ production via a scaled down Griess reaction method. At the beginning of incubation, samples for determination of cell density using flow cytometry were taken.

### Validation of the testing system with known SNIs and BNIs

Various NIs were utilized at a range of concentrations, expected to impose from slight to full inhibition of the activity of AOM strains. Different concentration levels were employed for the different reporter strains, based on their sensitivity, as depicted from earlier studies (33,37) [Table S2].

For AOB, 200 μL of the concentrated culture were added at each well, followed by addition of each inhibitor from stock solutions. Control cultures of ddH_2_O and DMSO were included as well. The final DMSO concentration of 0.1% v/v was not expected to impose significant inhibition on AOM (49,50). Controls and hydrophilic NIs (DMPP and MHPP) were initially diluted in sterile growth medium, before adding 20 μL to the wells. For the rest of the NIs, 0.2 μL were applied directly to the wells followed by the addition of 19.8 μL of growth medium. Triplicate cultures for each inhibitor x concentration combination were prepared, with each well representing one biological replication. Nitrite determination at different time points was used to determine the Ammonia Oxidation Inhibition percentage (AOI%) and the EC_50_ value.

For AOA, 200µL concentrated cultures were incubated with the addition of the respective NI or control in triplicates in 96-well plates for 3-5 hours. NI stocks in DMSO were diluted with filter-sterilized ddH_2_O to achieve a range of NI concentrations and not to exceed 1% (v/v) agent in the culture. Final DMSO concentrations in the assays did not exceed 0.02% (v/v), following the tolerance limit for the highly sensitive concentrated AOA cultures. Nitrite determination at a single time point was used to determine the Ammonia Oxidation Inhibition percentage (AOI%) and the EC_50_ value.

### Proof-of-concept of the fast-track testing system with wheat root exudates

Dried root exudates, derived as described before by Ghatak et al. (51), were resuspended in sterile ddH_2_O, and further diluted to standardize their concentration based on the original dry root weight [Table S3]. Inhibitory activity against the different AOM strains was determined at a single concentration level of 9.7 μg μL^-1^ [Table S3]. Control treatments with 6 μL of ddH_2_O were also included. The 96-well plates were incubated under optimum conditions for each of the AOM strain as described above. Nitrite determination was used to follow the inhibition of the AOM cells and determine the AOI% at one selected time – point, approximately 5 h for *N. viennensis* and *“Ca.* N. franklandianus*”*, 20 h for *N. multiformis*, and 22 h for *N. ureae*. Due to limited availability of root exudates, *N. communis*, the least sensitive AOB reporter strain, as derived from testing with pure BNIs and SNIs, was excluded from the screening process.

### Cell density determination

For AOB, DNA from collected samples was extracted using the NucleoSpin® Tissue kit (Macherey – Nagel GmbH & Co. KG, Düren, Germany), following the manufacturer’s protocol. The *amoA* gene was quantified via qPCR in a Biorad (Hercules, California, USA) CFX Real–Time PCR system, at 40 cycles, with the amoA-1F/amoA-2R set of primers (52) as described in Bachtsevani et al. (26). qPCR efficiency was > 80% and the R^2^ of the standard curve was > 0.98. For cell density determination, it was considered that the *amoA* gene is present in three copies in *N. multiformis* (53) and *N. ureae* (54), and in two copies in *N. communis* (55).

For AOA, cell abundance was quantified by flow cytometry as in Malits et al. (56). Samples were analysed using a FACSAria II (BD Biosciences, Franklin Lakes, New Jersey, USA) flow cytometer and AOA populations were discriminated and enumerated according to their specific signature in plots of 90° light scatter versus green DNA fluorescence.

### Data analyses

Modelling of the AOB routine cultures activity employed a single sigmoidal curve fit to nitrite production data obtained over consecutive generations, according to Caglar et al. (57). AOB maximum specific growth rate (μ_max_) was determined from the semi logarithmic plot of nitrite concentration against time, as performed in others (32,58). Activity rates of concentrated AOB cultures were calculated similarly, incorporating data from the entire incubation period.

EC_50_ values represent the concentrations that inhibit ammonia oxidation activity by 50% compared to the respective control treatment. Ammonia Oxidation Inhibition percentage (AOI%) refers to the proportional inhibition of AOM activity compared to the control. For AOB dose-response curves fitting a single sigmoidal pattern, normalized nitrite production data were used for EC_50_ determination, as described in Papadopoulou et al. (37). For linear or polynomial AOB responses, the slopes of the activity linear regression curves at each dose level were utilised to determine the AOI% for each inhibitor concentration. AOI% were plotted against the respective concentrations to estimate the EC_50_. For AOA data, AOI% values for one time-point were calculated as the proportional production of nitrite relative to the control, and EC_50_ values were determined as for the AOB. AOI% values for wheat root exudates were calculated as the proportional nitrite production compared to the control. For a more detailed description of the data analysis and resources used, refer to the Supplementary Material.

### Statistical tests and analyses

For data distribution validation, the Shapiro-Wilk normality test was conducted. Kruskal-Wallis test, and the Dunn’s post-hoc test with the Bonferroni p-value adjustment method was performed for comparing cell density means obtained from qPCR. Differences in the normalised nitrite values for the different SNI and BNI doses tested against AOA were evaluated with Kruskal-Wallis tests, and the Dunn’s post-hoc test with the Bonferroni p-value adjustment method. The heatmap of the BNI and SNI EC_50_ values was generated with hierarchical clustering and Euclidean distances for both column and row dendrograms. Comparisons of mean EC_50_ values per NI and AOM strain were performed utilising the respective means and standard errors and calculating the 95% confidence interval range. The RE AOI% values heatmap was generated with hierarchical clustering and Euclidean distances for both column and row dendrograms. Principal Component Analysis (PCA), to identify different inhibitory patterns, was conducted with scaled AOI% values. For a more comprehensive description of the statistical analyses, and the resources used, refer to the Supplementary Material.

## Results

### Fast – track system standardisation

Cell density and activity of concentrated AOM strain cultures were monitored in 96-well plates for consecutive tests and assessed relative to routine, non-concentrated cultures. For *N. multiformis*, cell harvesting significantly increased the cell density 5.6-fold compared to the logarithmic phase of the routine cultures (*p* < 0.001), and a further 2.6-fold increase was achieved with centrifugation (*p* < 0.05) [Table S4; Figure S1A]. The average cell density of the concentrated cultures per test well (3.57×10^7^ cells mL^-1^) did not significantly differ with that of the late logarithmic phase (*p* = 1) [Table S4]. Cell density trends were consistent across tests [Figure S1B]. The concentrated cultures displayed consistent activity, with nitrite production following a clear linear pattern (adjusted R^2^ = 0.9828, *p* < 0.001) [Figure S1C], with an activity rate of 0.113 ± 0.004, indicating a 2.5-fold increase in activity compared to the μ_max_ at the logarithmic phase of the routine cultures [Figure S4A and S4B; Table S5]. *N. ureae* concentrated cultures had, per test well, 9.43×10^7^ cells mL^-1^, an average cell density 2.4-fold higher than the late logarithmic growth phase of routine cultures (*p* < 0.001) [Table S4; Figure S2A], with consistent cell density trends across consecutive tests [Figure S2B]. Nitrite production followed a linear trend (adjusted R^2^ = 0.9945, *p* < 0.001) and the activity rate for the concentrated cultures was 0.160 ± 0.007, approximately 4.5 times higher than the μ_max_ at the logarithmic phase of the routine cultures [Figure S2C and S2D; Table S5]. *N.* communis concentrated cultures achieved average cell density of 1.54×10^5^ cells mL^-1^ per test well, a 4.9-fold higher cell density than routine cultures at the late logarithmic phase (*p* < 0.05) [Table S4; Figure S3A]. The slightly lower repeatability in the harvesting procedure [Figure S3B] did not affect the activity of the concentrated cultures [Figure S3C], which showed consistent nitrite production (adjusted R^2^ = 0.9943, *p* < 0.001) [Figure S3C], with an activity rate of 0.132 ± 0.002 [Figure S4E and S4F; Table S5], reflecting a 4-fold increase in activity compared to the μ_max_ at the logarithmic phase of the routine cultures.

The concentrated AOA cultures exhibited overall adjusted R^2^ values of 0.9811 (*p* < 0.001) for “*Ca.* N. franklandianus” [Figure S5A] and 0.9908 (*p* < 0.001) for *N. viennensis* [Figure S6A]. Strong congruence between the cell activity and the cell density curve was revealed for both strains [Figures S5B and S6B]. The concentrated cultures were initiated with similar cell densities, 1.28×10^7^ (± 1.79×10^6^) cells mL^-1^ for “*Ca.* N. franklandianus”, a 2.5-fold increase compared to the late logarithmic phase of the routine cultures [Table S4], and 8.70×10^7^ (± 1.12×10^5^) cells mL^-1^ for *N. viennensis*, approximately 6 times higher than the late logarithmic phase of the routine cultures [Table S4].

### Fast – track system validation with known SNIs and BNIs

All nitrapyrin doses inhibited AOB strains [Figures 2 and S7]. *N. ureae* was the most sensitive (EC_50_ = 0.18 ± 0.01 μM), followed by *N. multiformis* (EC_50_ 0.44 ± 0.03 μM) and *N. communis* (EC_50_ 0.88 ± 0.05 μM), with statistically different EC_50_ values [Figure 4]. AOA showed a weaker response [Figures 3 and S8], with “*Ca.* N. franklandianus” exhibiting an EC_50_ of 33.6 ± 4.1 μΜ [Figure 4]. DMPP showed a clear dose – response pattern for all AOM, with all applied doses inhibiting AOB [Figures 2 and S7]. *N. multiformis* (EC_50_ 0.48 ± 0.05 μM) was the most sensitive AOM, followed by *N. ureae* and *N. communis* [Figure 4]. DMPP was less effective on AOA, with statistically similar EC_50_ values of 1 847 ± 87 μM for *N. viennensis* and 1 890 ± 278 μM for “*Ca.* N. franklandianus” [Figure 4]. Ethoxyquin exhibited EC_50_ values of 2.18 ± 0.06 μM for *N. viennensis* and 6.37 ± 0.62 μM for “*Ca.* N. franklandianus” [Figure 4]. Amongst AOB, *N. ureae* showed the highest sensitivity (EC_50_ = 141.3 ± 43.2 μM) to ethoxyquin, while *N. multiformis* and *N. communis* required higher doses to achieve significant inhibition [Figures 2 and S7] (EC_50_ > 500 μM).

**Figure 2.**
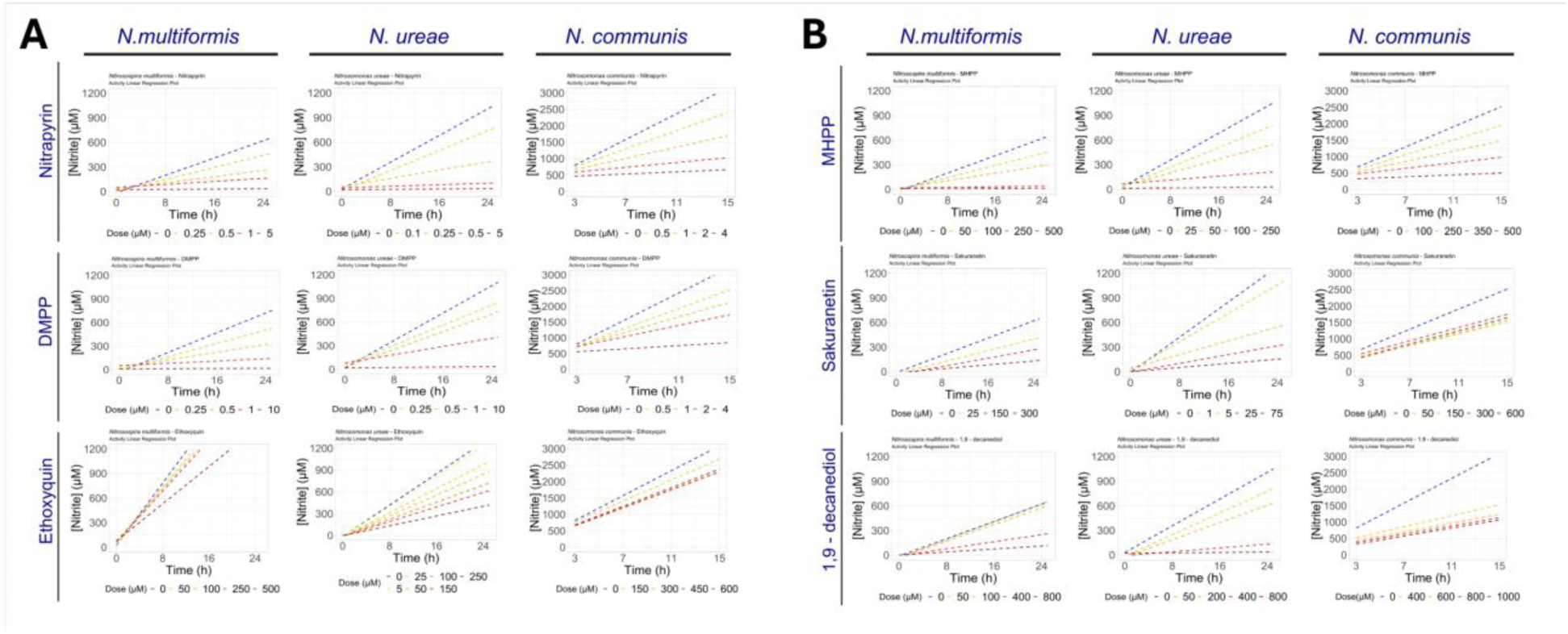
Validation of the AOB fast-track assay with pure SNIs and BNIs. The response of the AOB strains to the tested Biological Nitrification Inhibitors (BNIs) and Synthetic Nitrification Inhibitors (SNIs) is presented. Each of the six panels corresponds to one NI and is divided into three columns, each representing one AOB strain. For each strain and NI combination, the ammonia oxidation activity is illustrated through a linear regression plot, comparing the AOB response to different dose rates, alongside the respective control treatment. The dashed lines in each plot represent the linear regression of nitrite values (data points not shown) at each time point for each treatment. Calculations for the AOB response and EC_50_ values were based on data from multiple time points. Details on the modelling procedure are provided in the Materials and Methods section. Outputs of the modelling procedure are presented in the respective supplementary figure (Figure S7) in the Supplemental Material.

**Figure 3.**
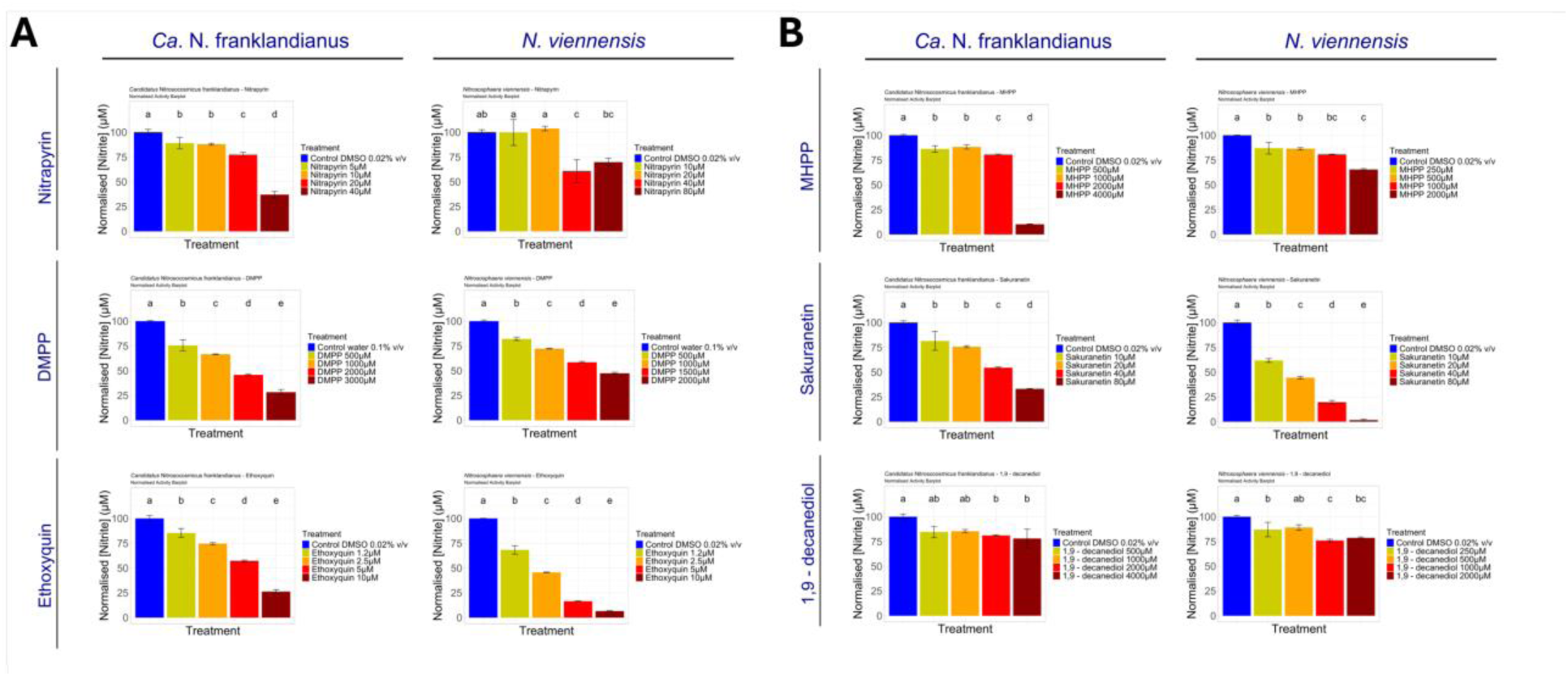
Validation of the AOA fast-track assay with pure SNIs and BNIs. The response of the AOA strains to the tested Biological Nitrification Inhibitors (BNIs) and Synthetic Nitrification Inhibitors (SNIs) is presented. Each of the six panels of the figure corresponds to one nitrification inhibitor and is composed of two columns. For each strain and NI combination, the normalized ammonia oxidation activity data are presented in a bar plot, comparing the AOA response to different dose rates, alongside the respective control treatment. The colours of the bars correspond to different dose rates, as indicated in the legend on the right. Lowercase letters indicate the grouping of treatments based on the Kruskal – Wallis test. Below the plot, a summary of the AOA response modelling, which led to the estimation of each EC_50_ value, is provided, utilizing a single model fit for the EC_50_ estimation. Calculations for the AOA response and EC_50_ values were based on data from a single time point. Detailed information about the modelling procedure can be found in the Materials and Methods section. Outputs of the modelling procedure are presented in the respective supplementary figure (Figure S8) in the Supplemental Material.

**Figure 4.**
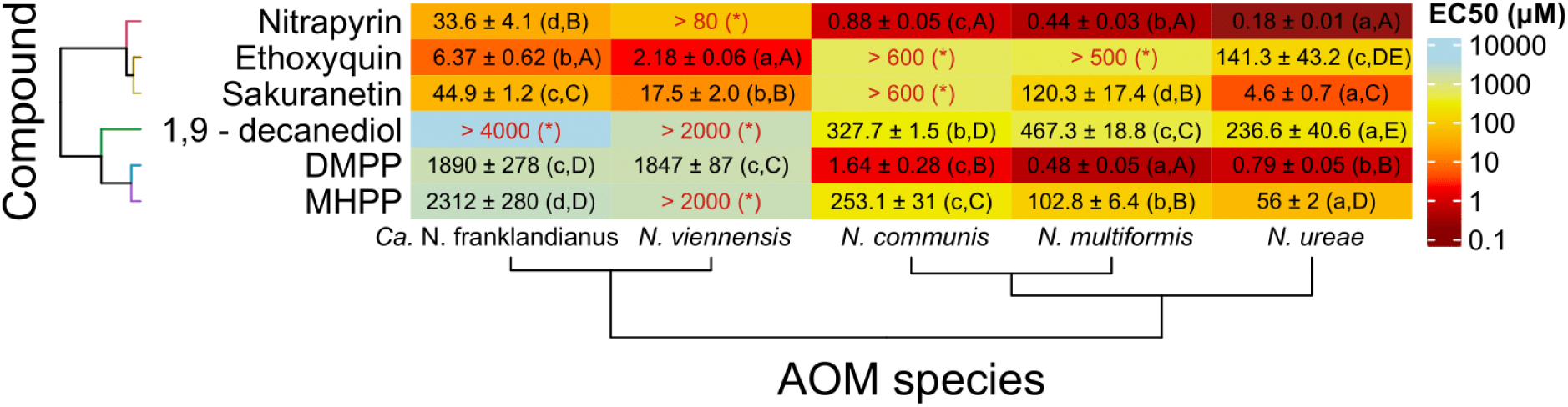
Comparison of NI inhibitory thresholds across the ammonia oxidizing microorganisms (AOM) reporter strains. A heatmap presenting the EC_50_ values for each nitrification inhibitor (NI) across the five reporter AOM strains used in the study, along with their respective standard errors. The colours of the boxes scale logarithmically according to the EC_50_ values, as indicted by the legend on the right. Dendrograms on the rows and columns were generated using hierarchical clustering of AOM strains and NIs based on Euclidean distances. Values denoted in red are indefinite EC_50_ values. Lowercase letters in parentheses indicate the grouping of EC_50_ values for each NI across strains, derived from comparison at a 95% confidence interval. Similarly, uppercase letters signify the grouping of EC_50_ values of all NIs per AOM strain, obtained from the same comparison. Asterisks (*) accompany indefinite values to indicate that they could not be reasonably compared to other values, while the colour of the respective boxes scales according to the legend based on the lowest limit value presented in each box.

MHPP exhibited dose – response patterns for all AOB [Figures 2 and S7], being most effective against *N. ureae* (EC_50_ = 56 ± 2 μM), while inhibition thresholds were significantly higher for *N. multiformis* (EC_50_ = 102.8 ± 6.4 μM) and *N. communis* (EC_50_ = 253.1 ± 31 μM) [Figure 4]. AOA showed low sensitivity to MHPP (EC_50_ > 2 000 μM) [Figure 4]. Sakuranetin exhibited higher inhibitory activity against *N. ureae* (EC_50_ 4.6 ± 0.7 μM) compared to *N. multiformis* (EC_50_ = 120.3 ± 17.4 μM), while *N. communis* did not show a clear dose-response (EC_50_ > 600 μM) [Figure 4]. AOA strains were sensitive to sakuranetin, with *N. viennensis* and “*Ca.* N. franklandianus” showing EC_50_ values of 17.5 ± 2 μM and 44.9 ± 1.2 μM respectively [Figure 4]. Lastly, 1,9-decanediol displayed the lowest efficacy amongst the NIs tested [Figures 3 and S8]. For the AOA strains, the highest concentrations tested failed to achieve > 50% inhibition [Figure 4], while AOB showed relatively higher sensitivity, with *N. ureae,* being the most sensitive strain (EC_50_ 236.6 ± 40.6 μM) [Figure 4].

Overall, nitrapyrin was the most potent NI against AOB, also active towards AOA, while ethoxyquin was the most effective against AOA [Figure 4]. DMPP, MHPP and 1,9-decanediol primarily targeted AOB, whereas the efficacy of sakuranetin was more strain-than domain-dependent. Notably, *N. ureae* was the most sensitive AOM strain against four of the six NIs tested (nitrapyrin, sakuranetin, 1,9-decanediol, and MHPP) [Figure 4].

Correlation analysis of EC_50_ values for *N. multiformis* against literature data yielded a high-quality linear regression [Figure S9A and S9B]. Excluding the ethoxyquin data, which deviated from the literature, significantly improved the correlation (R^2^ = 0.9907, slope = 1.09; data not shown), with no discrepancies observed for the other tested NIs [Figure S9C and S9D]. “*Ca.* N. franklandianus” inhibition thresholds were similar to literature values for sakuranetin and DMPP, and comparable for ethoxyquin [Figure S10C and S10D], showing good correlation and slope [Figure S10A and 10B]. For *N. viennensis*, EC_50_ values reported in the literature are limited, and correlation levels were of lower quality [Figure S11A and 11B], but comparable results were obtained for nitrapyrin and sakuranetin [Figure 11C and 11D].

### Fast-track system proof-of-concept with wheat root exudates

Hierarchical clustering revealed distinct inhibitory patterns among the root exudates, allocating them into different clusters [Figure 5]. Cluster A comprised two REs with high inhibition against all AOM strains except for *N. viennensis*, with RE15 being the most efficacious exudate tested. Cluster B included two REs that specifically inhibited *N. multiformis*, while two REs correlated with *N. ureae* inhibition were assigned to Cluster C. RE20, placed in cluster D, exhibited inhibition exclusively against “*Ca.* N. franklandianus”, whereas Cluster E contained five REs with no significant inhibitory potential. Four REs from Austria in Cluster F displayed notably high and specific inhibition against “*Ca.* N. franklandianus”. RE16, assigned to cluster G, selectively inhibited *N. viennensis*, while Cluster H was characterized by lower levels of inhibition across the different reporter strains [Figure 5]. Principal Component Analysis (PCA) accounted for 79.4% of the original variation [Figure S12], reaffirming RE15 as the most potent RE, and distinctly separating root exudates targeting AOB (RE5, RE13 and RE15) and AOA (RE2, RE6, RE9 and RE16) [Figure S12].

**Figure 5.**
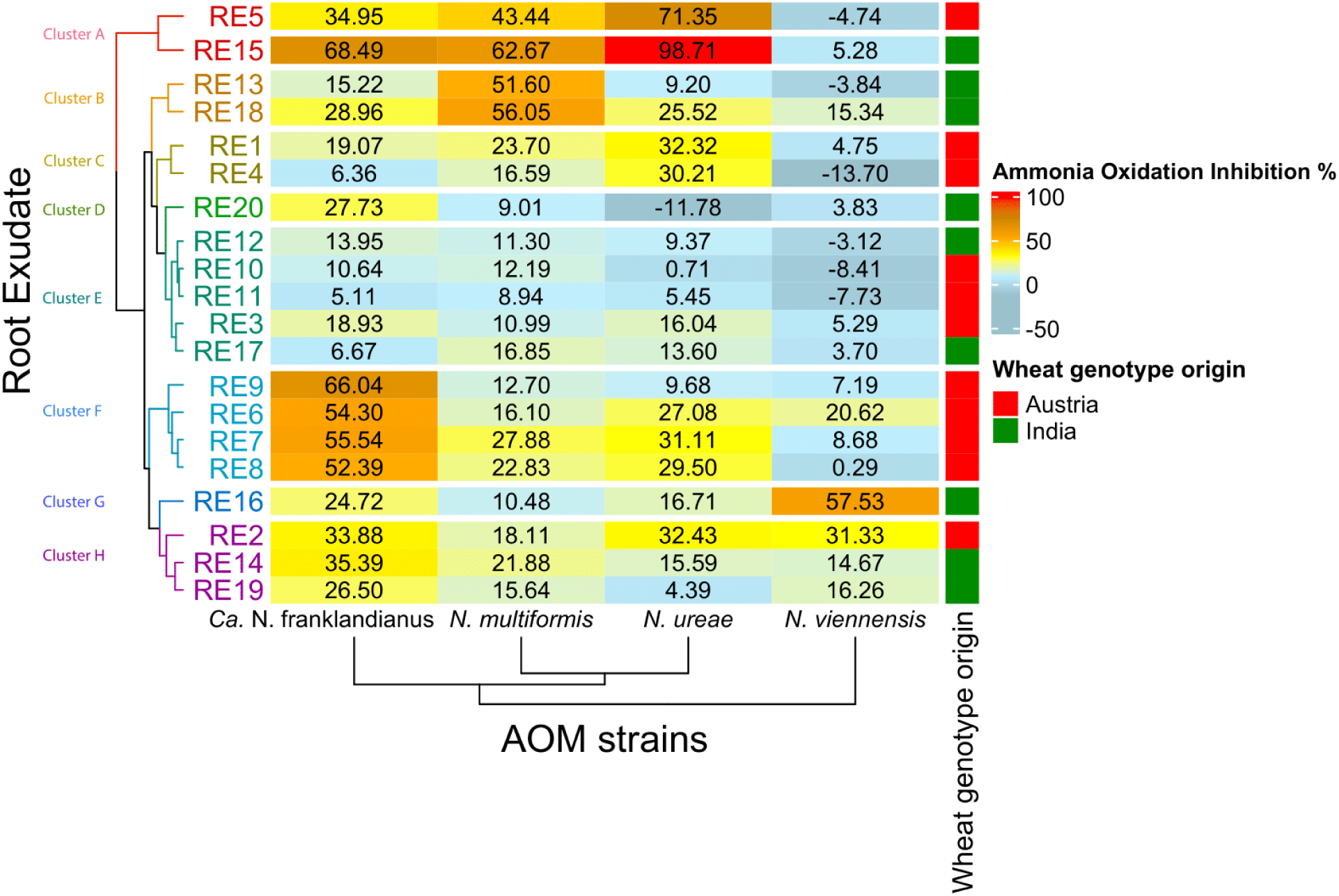
Comparison of root exudate inhibitory activity across the tested ammonia-oxidizing microorganism (AOM) strains. A heatmap showing the mean ammonia oxidation inhibition percentage values (AOI%) for each root exudate (RE) across four reporter AOM strains. The colours of the boxes scale according to the AOI% values, as indicted by the legend. Dendrograms on the rows and columns were generated using hierarchical clustering of AOM strains and root exudates based on Euclidean distances. The origin of the respective wheat genotypes is annotated on the right, with coloured boxes corresponding to different origin, as indicated in the additional legend. Due to low material availability, the two most sensitive AOB were employed to screen the wheat REs, and *N. communis* was excluded from the analysis.

## Discussion

In this study, we developed a high-throughput, fast-track screening assay for BNI activity. The turnaround time, ranging from 5 h for “*Ca.* N. frankandianus” and *N. viennensis* to 22 h for *N. ureae*, aligned with the studies of O’ Sullivan et al. (41) and Beeckman et al. (24). Shorter turnaround times, such as 30 min (38) and 2 h (31) rely on highly dense AOM cultures, which may underestimate the true NI potential, due to elevated metabolic activity, the presence of multiple molecular targets and the use of indirect endpoints (i.e., bioluminescence) instead of directly measuring nitrification activity (38). According to the "inoculum effect" (59,60), similar to antibiotics, the efficacy of NIs directly inhibiting ammonia monooxygenase (AMO) or other nitrification-related enzymes would appear reduced in highly dense cultures. For example, nitrapyrin, oxidized by AMO into 6-chloropicolinic acid (61), may be affected by the increased AMO concentration in dense cultures. Additionally, compounds such as 6-chloropicolinic acid, which exhibit up to 8 h delay in responses (62), may appear ineffective in assays employing shorter timescales, due to repressed BNI uptake and activity. BNI lipophilicity and polarity influence transfer across membranes and within intracellular spaces, both of which can differ between AOA and AOB (25). Longer incubation times employed here ensure capture of the full cellular response and a more precise reflection of the BNI potential in AOM reporter strains.

To advance the ecological relevance and phylogenetic diversity compared to previous efforts (24,31,38,41), we employed three phylogenetically diverse, soil-representative AOB strains. *Nitrosospira*, the most abundant AOB in soil (3), are crucial ammonia oxidisers in fertilized soils (63,64) and *N. multiformis* (*Nitrosospira* Cluster 3), highly adaptive in agricultural soils (53,65), has been utilised as a model AOB in several previous studies (32,33,36,37). We also included *N. ureae* (*Nitrosomonas* Cluster 6a) and *N. communis* (*Nitrosomonas* Cluster 8) (66), replacing the less soil-prevalent *Nitrosomonas* representative *N. europaea.* Regarding AOA, “*Candidatus* Nitrosocosmicus franklandianus” (Clade NS-ζ-2) and *N. viennensis* (Clade NS-α- 3.2.1) (67) represent distinct soil-abundant clades. *N. viennensis* is one of the best studied soil AOA, with significant potential for N_2_O emission (17), utilized before to assess SNIs (36) and BNIs (32), including fast-track approaches (24). “*Ca.* Nitrosocosmicus franklandianus” represents a clade exhibiting high autotrophic growth in environmental samples (67), with phylogeny and classification distinct to *Nitrososphaera* (68,69). Notably, “*Ca.* Nitrosocosmicus” members, possessing a unique manganese catalase (MnKat) gene, have emerged as key nitrifiers in the rhizosphere of important crops, indicating a previously underexplored AOA-plant interaction (70). Overall, our fast-track assay, incorporating five AOM reporter strains from both microbial domains driving ammonia oxidation in soil and having soil-derived cultivable representatives, offers a more robust assessment of the differential responses of AOB and AOA, better reflecting the diversity of soil AOM communities.

AOM responses to pure NIs reflected the different NI efficacy and the differences in AOM sensitivity (32,33,37). Notably, these are the first reported inhibition thresholds for *N. communis* and *N. ureae*. The latter was more sensitive than *N. multiformis*, while *N. communis* was the least sensitive AOB strain tested. Nitrapyrin was highly potent against AOB, with *N. multiformis* EC_50_ values aligning with literature (37). *N. ureae* and *N. communis* were both more sensitive to nitrapyrin than *N. europaea,* for which reported EC_50_ values span from 2.1 ± 0.4 μM (37) to even higher (71), reinforcing the importance of prioritizing more ecologically relevant strains in such experimental schemes.

DMPP exhibited comparable to nitrapyrin inhibitory activity on AOB, with lower efficacy against AOA, in accord with previous studies (37,72). Inhibition thresholds for *N. multiformis* and “*Ca.* N. franklandianus” matched earlier studies (37), while *N. viennensis* exhibited a similar EC_50_ value to “*Ca.* N. franklandianus”. DCD, another weak AOA inhibitor, also produced comparable threshold values between *N. viennensis* and “*Candidatus* Nitrosocosmicus. agrestis” (36,73). The specificity of ethoxyquin towards AOA (37) was reaffirmed in this study, as the estimated EC_50_ for “*Ca.* N. franklandianus” was comparable to earlier findings (EC_50_ 1.4 ± 0.3 μM vs 6.37 ± 0.62 μM in this study) (37), and a similar value was obtained for *N. viennensis*. *N. multiformis* and *N. communis* were marginally inhibited, consistent with earlier findings for *N. multiformis* in batch cultures (EC_50_ = 214.8 ± 39.6 μM) (37).

MHPP, a BNI with strain-dependent inhibitory patterns (32,33), produced an EC_50_ value of 102.8 ± 6.4 μM for *N. multiformis*, agreeing with literature (33). Both AOA strains required much higher concentrations (∼ 2 000 μM) for inhibition. For *N. viennensis*, approximately 30% inhibition was observed at 2 000 μΜ, suggesting an EC_50_ higher than that of “*Ca.* N. franklandianus” (2 312 ± 280 μM), in agreement with previous findings (32). Sakuranetin exhibited no clear selectivity towards either AOB or AOA. *N. multiformis* had an EC_50_ value (120.3 ± 17.4 μM) closely matching values from Kolovou et al. (144.3 ± 19 μM) (33) and Kaur-Bhambra et al. (141.1 ± 0.6 μM) (32). The EC_50_ value for “*Ca.* N. franklandianus” (44.94 ± 1.15 μM) aligns with Kaur-Bhambra et al. (61 ± 10 μM) (32), though *N. viennensis* showed higher sensitivity. 1,9-decanediol was the weakest NI tested, showing comparatively higher activity towards AOB. *N. ureae* was the most sensitive, followed by *N. multiformis* and *N. communis*, the former showing EC_50_ value (467.3 ± 18.8 μM) aligning with earlier studies (428.1 ± 2.47 μM) (32). AOA inhibition thresholds exceeded the highest tested concentrations (> 2 000 μM) compared to literature values of 418.3 ± 19.2 μM and 211.7 ± 0.9 μM for *N. viennensis* and “*Ca.* N. franklandianus”, respectively (32).

Overall, *N. multiformis* thresholds were congruent with literature, whereas the two AOA strains showed discrepancies, likely due to the shorter exposure period in the fast-track assay, compared to the prolonged exposure in liquid culture assays, which may amplify weak NI potency. Variations were less evident for the potent AOA inhibitors such as ethoxyquin and nitrapyrin, which do not require extended exposure period to exert their toxicity. Conversely, for weaker NIs like 1,9-decanediol, longer testing periods likely allow for increased uptake, resulting in higher potency against AOA. Additionally, AOA biofilm production, significantly enhanced in dense cultures, could further reduce NI efficacy. Members of both the clade of *N. viennensis* (NS-α) (74) and the clade of “*Ca.* N. franklandianus” (NS-ζ) (75–77) have been found to form aggregates and biofilm-like structures, potentially limiting NI uptake. Moreover, the distinct cell wall features of NS-ζ clade members (69) and the S-layer of NS-α members, could impact NH_4_^+^ accumulation (78), further influencing NI activity. Given these physiological differences, variations in sensitivity between *N. viennensis* and “*Ca.* N. franklandianus” are expected.

Different inhibitory patterns were observed for the wheat root exudates, as a result of differences in AOM species sensitivity and root exudate chemical composition which often exhibits genotype-dependent patterns (79). Selective activity was noted against all four AOM strains. Notably, responses from *N. viennensis* did not cluster with its AOA counterpart [Figure 5], most probably due to the presence of different active molecules with variable inhibitory activity towards the tested AOA, as well as the differences in cell surface and growth mode between the two AOA (69). Differences in BNI activity have been reported before in root extracts from various maize (80), guinea grass (35), and rice cultivars (31), while geographical origin of wheat genotypes has also been shown to affect root exudate composition (81), though this was not evident in our study. Overall, the selective inhibition of specific AOM strains by certain root exudates, as observed in this study, likely reflects the adaptation of specific genotypes to their native soil environment, including local microbial diversity (82), soil type (83) and pH (83).

Overall, in this study, we developed a fast-track, high-throughput assay for BNI activity screening. Validated with known NIs and plant root extracts, the assay demonstrates robust applicability for large-scale screening of both pure compounds and plant-derived extracts. This system advances similar previous testing platforms by incorporating a broader range of more ecologically relevant, phylogenetically distinct AOB and AOA strains, laying the foundation for the systematic, high-throughput discovery of BNI plant genotypes and compounds in relevant contexts. When combined with more realistic soil-based testing systems, our assay holds promise for early discovery of novel BNIs and BNI-producing plants, addressing critical agricultural and environmental issues, such as NUE loss, nitrite pollution and greenhouse gas emissions contributing to climate change. Future improvements, such as further expanding the diversity of tested strains or refining the ability of the system to mimic natural soil environments, could further upgrade its ecological relevance.

## Acknowledgements

This work was funded by the Grantham Foundation as part of the project ‘Pipeline for Development and Commercialization of Biological Nitrification Inhibitors to Mitigate GHG Emissions from Cultivated Soils.’. Alexandros Kanellopoulos was partially funded by the MSc program “HOSMIC – Host-Microbe Interactions”. We thank Dr. Logan Hodgskiss and Mr. Maximilian Dreer (Archaea Biology and Ecogenomics Unit, Department of Functional and Evolutionary Ecology, Faculty of Life Sciences, University of Vienna) for assistance with the experimental setup of the AOA screening method and discussion of the results.

## Supplemental material

### Materials and Methods

#### Data analyses

Data processing and visualisation were conducted using the RStudio software (R Core Team 2020). Plot generation utilized either the “ggplot2” package v3.4.2 (Wickham, 2016) or base R functions. In the duration of the analyses, the R packages “readxl” v1.4.3 (Wickham and Bryan 2023), “dplyr” v1.1.4 (Wickham et al. 2023) and “tibble” v3.2.1 (Müller and Wickham 2023) have been used for general data processing purposes, and the “ggplot2” v3.5.0 (Wickham 2016), “ggpubr” v0.6.0 (Kassambara 2023a) and “patchwork” v1.2.0 (Pedersen, 2024) R packages have been used for the generation of figures.

Modelling of the activity of AOB routine cultures was performed by fitting a single sigmoidal curve to nitrite data obtained over consecutive generations. The modelling employed the “sicegar” package v0.2.4 (Caglar et al. 2018), using the equation that follows in (1), according to Caglar et al. (2018). AOB μ_max_ was determined from the semi logarithmic plot of nitrite concentration against time, as performed in others (Powell and Prosser 1986; Kaur – Bhambra et al. 2022). Activity rates of AOB concentrated cultures were calculated using the same approach, utilizing data from the entire incubation period.

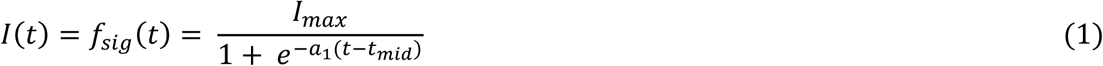

EC_50_ values represent the concentrations that inhibit ammonia oxidation activity by 50% compared to the respective control treatments. Ammonia Oxidation Inhibition percentage (AOI%) for pure NIs refers to the proportional inhibition of AOM activity compared to the control. For dose-response curves fitting a traditional single sigmoidal pattern, normalized nitrite production data were used with the “drc” v3.0.1 package (Ritz et al. 2016) to determine EC50 values, as described in Papadopoulou et al. 2020. EC_50_ values were generated by the algorithm for each time point, and an average value ± the standard error was calculated from those with a satisfactory p – value and R^2^ ≥ 0.95. For dose schemes that produced a linear or polynomial AOB response, the slopes of the AOM activity linear regression curves at each dose level were utilised to determine the AOI% for each inhibitor concentration, with calculations performed as shown in (2). In this case, the AOI% values were plotted against the respective concentrations. If a satisfactory linear regression (adjusted R^2^ > 0.9, p – value < 0.05) could be produced with at least 4 out of 5 data points, the EC_50_ concentration was calculated from the equation of the best linear regression, for y = 50. A similar approach was utilized by Kaur – Bhambra et al. (2022) with μ_max_ replacing the AOI%. The standard error of the EC_50_ value was considered to coincide with the standard error produced for the unknown predictor value by the *calibrate ()* function of the “investr” v1.4.2 R package (Greenwell and Schubert Kabban 2014) at a 95% confidence level, when the observed variable was set to 50. If no satisfactory linear relationship was observed, a third-degree polynomial fit was utilized instead, with the EC_50_ calculated from the equation of the curve for y = 50. The standard error was estimated through the Delta Method. Briefly, the first derivative of the 3^rd^ degree polynomial equation at the root for y = 50 was calculated and the sensitivity of the polynomial function with respect to its coefficients at the root was computed, in order to calculate variance propagation and deduct the standard error.

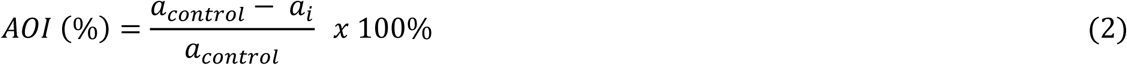

For AOA data, where one time – point was utilised, AOI% values calculated using the equations in (3-5) were plotted against the respective concentrations and a similar approach utilising a linear, a 3^rd^ degree polynomial or a log_10_ fit, with the model decided based on the best fit. Standard errors were estimated as described above.

AOI% for wheat root exudates refers to the proportional inhibition of AOM activity compared to the control. It was calculated for each treatment as follows in (5), after normalising all nitrite production values (3-4).

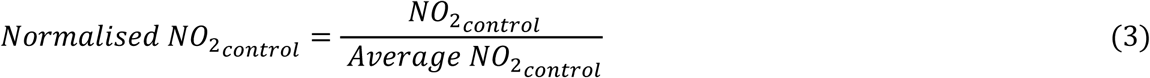

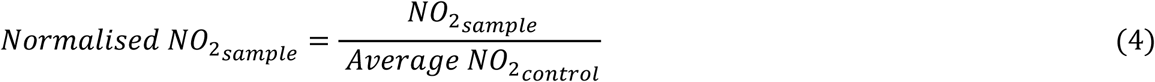

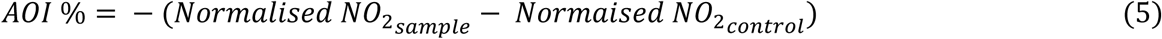

#### Statistical tests and analyses

For data distribution validation, the Shapiro-Wilk normality test was conducted, supplemented by the respective histogram and q-q plot using the “stats” base package in the R Software. Kruskal-Wallis test and the Dunn’s post-hoc test with the Bonferroni p-value adjustment method was performed using the “agricolae” package v1.3.7 (de Mendiburu, 2023) and the “rstatix” v0.7.2 (Kassambara 2023b) R packages for comparing cell density means obtained from qPCR. Differences in the normalised nitrite values for the different SNI and BNI doses tested against AOA were evaluated with Kruskal-Wallis tests, and the Dunn’s post-hoc test with the Bonferroni p-value adjustment method, employing the “agricolae” package v1.3.7 (de Mendiburu, 2023) and the “rstatix” v0.7.2 (Kassambara 2023b) R packages. The heatmap of the BNI and SNI EC_50_ values were generated with the utilisation of the “ComplexHeatmap” v2.14.0 (Gu et al. 2016) and “dendextend” v1.17.1 (Galili 2015) R packages, with both column and row dendrograms based on hierarchical clustering and Euclidean distances. Comparisons of these mean EC_50_ values per NI and AOM strain were performed utilising the respective means and standard errors and calculating the 95% confidence interval range based on the equation in (6).

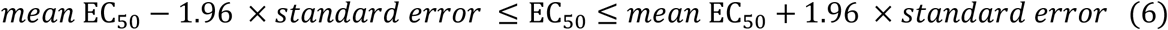

The heatmap of the RE AOI% values was generated with the utilisation of the “ComplexHeatmap” v2.14.0 (Gu 2016) and “dendextend” v1.17.1 (Galili 2015) R packages, with both column and row dendrograms based on hierarchical clustering and Euclidean distances. Finally, Principal Component Analysis (PCA) to identify different inhibitory patterns, was conducted after AOI% values were scaled using the “vegan” v2.6.4 R package (Oksanen et al. 2022) and the plot was generated using the “factoextra” v1.0.7 R package (Kassambara and Mundt 2020).

## Supplementary Tables

**Supplementary Table S1.** Summary of the provider and purity of each compound used in the study.

| Compound | Provider | Purity |
| --- | --- | --- |
| **DMPP** | Cayman Chemical (1180 East Ellsworth Road, Ann Arbor, Michigan, 48108, USA) | ≥ 98% |
| **Nitrapyrin** | Sigma – Aldrich (Chemie GmbH, Eschenstr. 5, 82024, Taufkirchen, Germany) | ≥ 98% |
| **Ethoxyquin** | Sigma – Aldrich (Chemie GmbH, Eschenstr. 5, 82024, Taufkirchen, Germany) | 96.9% |
| **MHPP** | BLDpharm (No.1 Wangdong Middle Road, Building 5, Songjiang District, Shanghai 201601, China) | 99.21% |
| **Sakuranetin** | Syngenta Crop Protection AG (Rosentalstr. 67, 4058 Basel, Switzerland) | > 95% |
| **1,9-decanediol** | WuXi AppTec (288 Fute Zhong Road, Shanghai, 200131, China) | 95% |

Abbreviations: DMPP – 3,4-dimethylpyrazole (phosphate), MHPP - methyl 3-(4-hydroxyphenyl) propionate

**Supplementary Table S2.** Overview of the concentrations of the nitrification inhibitors used for the validation of the fast-track testing system.

|  | Compound | Tested concentrations (μM) |  |  |  |
| --- | --- | --- | --- | --- | --- |
| *SNIs* | DMPP | 0.25 | 0.5 | 1 | 10 |
|  | Nitrapyrin | 0.25 | 0.5 | 1 | 5 |
|  | Ethoxyquin | 50 | 100 | 250 | 500 |
| *BNIs* | MHPP | 50 | 100 | 250 | 500 |
|  | Sakuranetin | 25 | 75 | 150 | 300 |
|  | 1,9-decanediol | 50 | 100 | 400 | 800 |

***Nitrosomonas ureae***
|  | Compound | Tested concentrations (μM) |  |  |  |  |  |
| --- | --- | --- | --- | --- | --- | --- | --- |
| *SNIs* | DMPP | 0.25 | 0.5 | 1 | 10 |  |  |
|  | Nitrapyrin | 0.1 | 0.25 | 0.5 | 5 |  |  |
|  | Ethoxyquin | 5 | 25 | 50 | 100 | 150 | 250 |
| *BNIs* | MHPP | 25 | 50 | 100 | 250 |  |  |
|  | Sakuranetin | 1 | 5 | 25 | 75 |  |  |
|  | 1,9-decanediol | 50 | 100 | 400 | 800 |  |  |

***Nitrosomonas communis***
|  | Compound | Tested concentrations (μM) |  |  |  |  |
| --- | --- | --- | --- | --- | --- | --- |
| *SNIs* | DMPP | 0.5 | 1 | 2 | 4 |  |
|  | Nitrapyrin | 0.5 | 1 | 2 | 4 |  |
|  | Ethoxyquin | 50 | 150 | 300 | 450 | 600 |
| *BNIs* | MHPP | 100 | 250 | 350 | 500 |  |
|  | Sakuranetin | 25 | 50 | 150 | 300 | 600 |
|  | 1,9-decanediol | 400 | 600 | 800 | 1000 |  |

***“Candidatus Nitrosocosmicus franklandianus”***
|  | Compound | Tested concentrations (μM) |  |  |  |
| --- | --- | --- | --- | --- | --- |
| *SNIs* | DMPP | 500 | 1000 | 2000 | 3000 |
|  | Nitrapyrin | 5 | 10 | 20 | 40 |
|  | Ethoxyquin | 1.2 | 2.5 | 5 | 10 |
| *BNIs* | MHPP | 500 | 1000 | 2000 | 4000 |
|  | Sakuranetin | 10 | 20 | 40 | 80 |
|  | 1,9-decanediol | 500 | 1000 | 2000 | 4000 |

***Nitrososphaera viennensis***
|  | Compound | Tested concentrations (μM) |  |  |  |
| --- | --- | --- | --- | --- | --- |
| *SNIs* | DMPP | 500 | 1000 | 1500 | 2000 |
|  | Nitrapyrin | 10 | 20 | 40 | 80 |
|  | Ethoxyquin | 1.2 | 2.5 | 5 | 10 |
| *BNIs* | MHPP | 250 | 500 | 1000 | 2000 |
|  | Sakuranetin | 10 | 20 | 40 | 80 |
|  | 1,9-decanediol | 250 | 500 | 1000 | 2000 |

Abbreviations: SNIs – Synthetic Nitrification Inhibitors, BNIs – Biological Nitrification Inhibitors, DMPP – 3,4-dimethylpyrazole (phosphate), MHPP - methyl 3-(4-hydroxyphenyl) propionate

**Supplementary Table S3.** A list of the wheat root exudates used for proof of concept and their concentrations applied in the study.

| **Sample number** | **Genotype** | **Place** | **Dried root weight (g)** | **Final Diluted Concentration ($\mu\text{g } \mu\text{L}^{-1}$)** | **Volume Added ($\mu\text{L}$)** | **Final Dose ($\mu\text{g } \mu\text{L}^{-1}$)** |
| --- | --- | --- | --- | --- | --- | --- |
| **RE1** | Adrianus | Austria | 1.14 | 333.33 | 6 | 9.7 |
| **RE2** | Alessio | Austria | 0.73 | 333.33 | 6 | 9.7 |
| **RE3** | Aristeus (SZD 056-112) | Austria | 1.23 | 333.33 | 6 | 9.7 |
| **RE4** | Arnold | Austria | 0.81 | 333.33 | 6 | 9.7 |
| **RE5** | Asli | Austria | 0.53 | 333.33 | 6 | 9.7 |
| **RE6** | Aurelius | Austria | 0.67 | 333.33 | 6 | 9.7 |
| **RE7** | Axaro | Austria | 0.81 | 333.33 | 6 | 9.7 |
| **RE8** | Balaton | Austria | 0.73 | 333.33 | 6 | 9.7 |
| **RE9** | Mandarin | Austria | 0.64 | 333.33 | 6 | 9.7 |
| **RE10** | Tillsano | Austria | 0.52 | 333.33 | 6 | 9.7 |
| **RE11** | Tobias | Austria | 0.82 | 333.33 | 6 | 9.7 |
| **RE12** | IW-D-3 | India | 0.36 | 333.33 | 6 | 9.7 |
| **RE13** | HUW669 | India | 0.36 | 333.33 | 6 | 9.7 |
| **RE14** | IW-D-2 | India | 0.32 | 333.33 | 6 | 9.7 |
| **RE15** | C.R4 | India | 0.52 | 333.33 | 6 | 9.7 |
| **RE16** | DBW187 | India | 0.48 | 333.33 | 6 | 9.7 |
| **RE17** | SR.S4 | India | 0.3 | 333.33 | 6 | 9.7 |
| **RE18** | RAJ3765 | India | 0.43 | 333.33 | 6 | 9.7 |
| **RE19** | IW-D-1 | India | 0.48 | 333.33 | 6 | 9.7 |
| **RE20** | IW-D-59 | India | 0.56 | 333.33 | 6 | 9.7 |

**Supplementary Table S4.** Overview of cell densities at different points of the AOM routine cultures and harvesting protocol. For AOB, data are available for two extra steps referring to the cell harvesting procedure. Values represent the average cells mL⁻¹ of three consecutive tests (± standard deviation) (SD). Values designated by different lowercase letters belong to groups statistically different at the 5% level, as determined by the Kruskal-Wallis test. “Routine culture initial” corresponds to the starting point of the routine cultures. “Routine culture final” corresponds to the late logarithmic growth phase of the routine cultures, just before AOB cell harvesting. The term “Inocula after cell harvesting” refers to the “After cell harvesting” step in Figures S1-S3.

| Step | Cell density (cells mL<sup>-1</sup>) (± 1SD) |
| --- | --- |
| Routine culture initial | 2.86×10<sup>5</sup> (± 1.45×10<sup>5</sup>) d |
| Routine culture final | 3.02×10<sup>7</sup> (± 9.55×10<sup>6</sup>) c |
| Inocula after cell harvesting | 1.69×10<sup>8</sup> (± 7.09×10<sup>7</sup>) b |
| Merged inoculum | 4.35×10<sup>8</sup> (± 4.17×10<sup>7</sup>) a |
| Concentrated culture initial | 3.57×10<sup>7</sup> (± 2.10×10<sup>6</sup>) c |

*Nitrosomonas ureae*
| Step | Cell density (cells mL<sup>-1</sup>) (± 1SD) |
| --- | --- |
| Routine culture initial | 3.62×10<sup>5</sup> (± 1.76×10<sup>5</sup>) e |
| Routine culture final | 3.95×10<sup>7</sup> (± 4.41×10<sup>6</sup>) d |
| Inocula after cell harvesting | 5.12×10<sup>8</sup> (± 1.15×10<sup>8</sup>) b |
| Merged inoculum | 1.62×10<sup>8</sup> (± 1.75×10<sup>8</sup>) a |
| Concentrated culture initial | 9.43×10<sup>7</sup> (± 1.16×10<sup>7</sup>) c |

*Nitrosomonas communis*
| Step | Cell density (cells mL<sup>-1</sup>) (± 1SD) |
| --- | --- |
| Routine culture initial | 1.14×10<sup>3</sup> (± 6.02×10<sup>2</sup>) d |
| Routine culture final | 3.96×10<sup>4</sup> (± 2.35×10<sup>4</sup>) c |
| Inocula after cell harvesting | 1.36×10<sup>6</sup> (± 4.46×10<sup>5</sup>) a |
| Merged inoculum | 2.79×10<sup>6</sup> (± 1.24×10<sup>6</sup>) a |
| Concentrated culture initial | 1.54×10<sup>5</sup> (± 8.79×10<sup>4</sup>) b |

| ***“Candidatus Nitrosocosmicus franklandianus”*** |  |
| --- | --- |
| **Step** | **Cell density (cells mL<sup>-1</sup>) (± 1SD)** |
| Routine culture initial | 5.15×10<sup>4</sup> (± 6.20×10<sup>3</sup>) c |
| Routine culture final | 5.15×10<sup>6</sup> (± 5.07×10<sup>5</sup>) b |
| Concentrated culture initial | 1.28×10<sup>7</sup> (± 1.79×10<sup>6</sup>) a |
| ***Nitrososphaera viennensis*** |  |
| **Step** | **Cell concentration (cells mL<sup>-1</sup>) (± 1SD)** |
| Routine culture initial | 7.20×10<sup>3</sup> (± 5.27×10<sup>2</sup>) c |
| Routine culture final | 1.44×10<sup>7</sup> (± 1.05×10<sup>6</sup>) b |
| Concentrated culture initial | 8.70×10<sup>7</sup> (± 1.12×10<sup>5</sup>) a |

**Supplementary Table 5.**
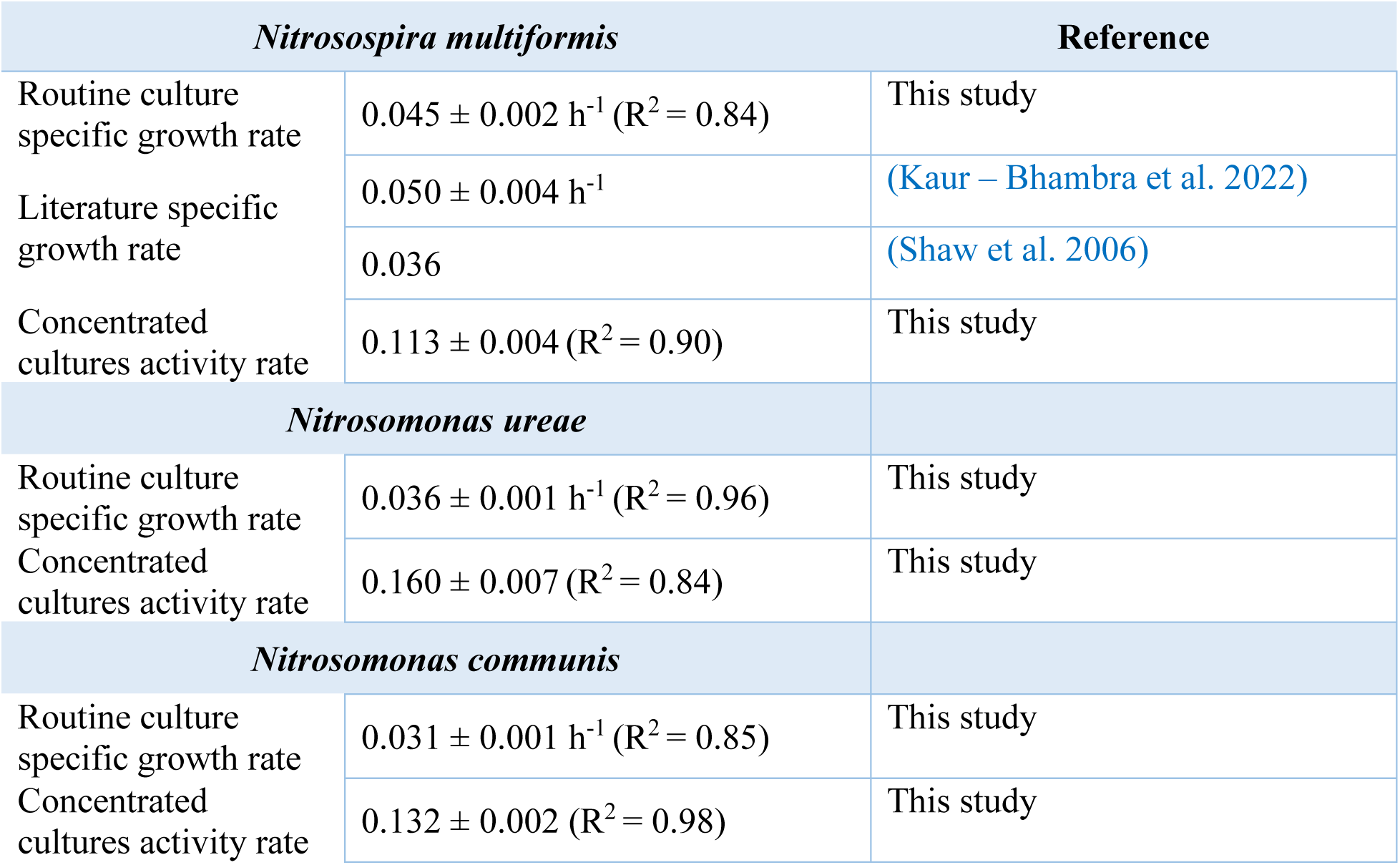
Overview of AOB specific growth rates and concentrated cultures activity rates. μ_max_ and activity rates are presented (± standard error). R^2^ values refer to the linear regressions utilized for μ_max_ and activity rates (shown in Figure S4).

| *Nitrosospira multiformis* |  | Reference |
| --- | --- | --- |
| Routine culture specific growth rate | $0.045 \pm 0.002 \text{ h}^{-1}$ ($R^2 = 0.84$) | This study |
| Literature specific growth rate | $0.050 \pm 0.004 \text{ h}^{-1}$ | (Kaur – Bhambra et al. 2022) |
|  | 0.036 | (Shaw et al. 2006) |
| Concentrated cultures activity rate | $0.113 \pm 0.004$ ($R^2 = 0.90$) | This study |
| *Nitrosomonas ureae* |  |  |
| Routine culture specific growth rate | $0.036 \pm 0.001 \text{ h}^{-1}$ ($R^2 = 0.96$) | This study |
| Concentrated cultures activity rate | $0.160 \pm 0.007$ ($R^2 = 0.84$) | This study |
| *Nitrosomonas communis* |  |  |
| Routine culture specific growth rate | $0.031 \pm 0.001 \text{ h}^{-1}$ ($R^2 = 0.85$) | This study |
| Concentrated cultures activity rate | $0.132 \pm 0.002$ ($R^2 = 0.98$) | This study |

## Supplementary Figures

**Figure S1.**
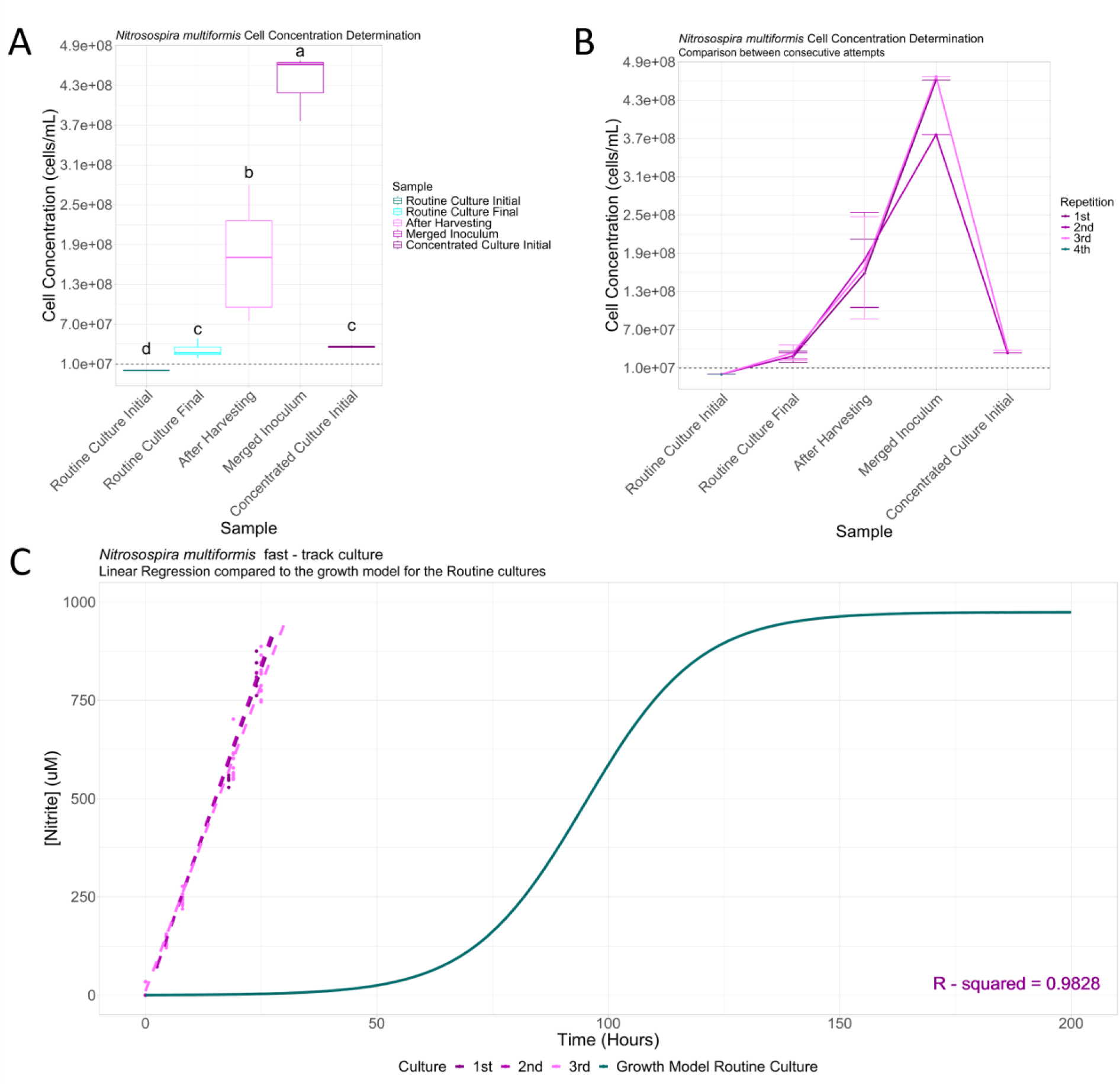
Nitrosospira multiformis fast – track assay protocol standardisation. (A) Box plot of pooled qPCR results for the different points of the cell harvesting procedure. The y – axis values correspond to cell concentration, while the five different points of the procedure are plotted on the x – axis. The black dashed line represents the limit below which the box plot is not to scale. Lowercase letters above the boxes indicate the grouping according to the Kruskal – Wallis test. (B) Line plots of qPCR results between consecutive tests of the cell harvesting procedure. Different tests are represented by different colours according to the legend. The coloured error bars represent the range between -1 and +1 standard deviation from the mean value. The black dashed line indicates the limit below which the line plot is not to scale. (C) Activity curve of N. multiformis fast – track culture (magenta lines) compared to the activity model curve of the respective routine cultures (blue line). The points represent individual nitrite values from the biological replicates at each time-point of each fast – track culture. The R^2^ value refers to the linear regression of the pooled fast – track culture data.

**Figure S2.**
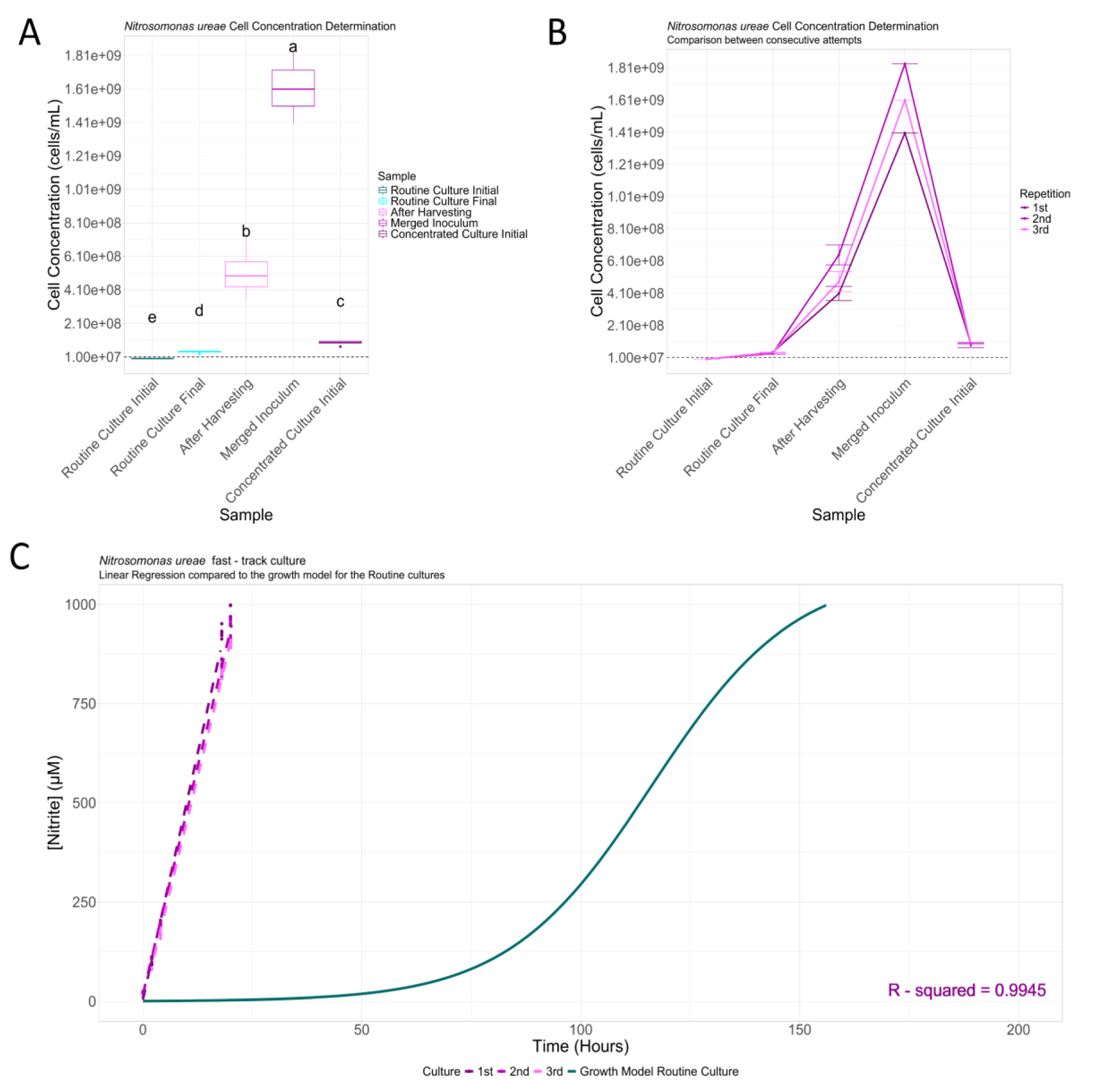
Nitrosomonas ureae fast – track assay protocol standardisation. (A) Box plot of pooled qPCR results for the different points of the cell harvesting procedure. The y – axis values correspond to cell concentration, while the five different points of the procedure are plotted on the x – axis. The black dashed line represents the limit below which the box plot is not to scale. Lowercase letters above the boxes indicate the grouping according to the Kruskal – Wallis test. (B) Line plots of qPCR results between consecutive tests of the cell harvesting procedure. Different tests are represented by different colours according to the legend. The coloured error bars represent the range between -1 and +1 standard deviation from the mean value. The black dashed line indicates the limit below which the line plot is not to scale. (C) Activity curve of N. ureae fast – track culture (magenta lines) compared to the activity model curve of the respective routine cultures (blue line). The points represent individual nitrite values from the biological replicates at each time-point of each fast – track culture. The R^2^ value refers to the linear regression of the pooled fast – track culture data.

**Figure S3.**
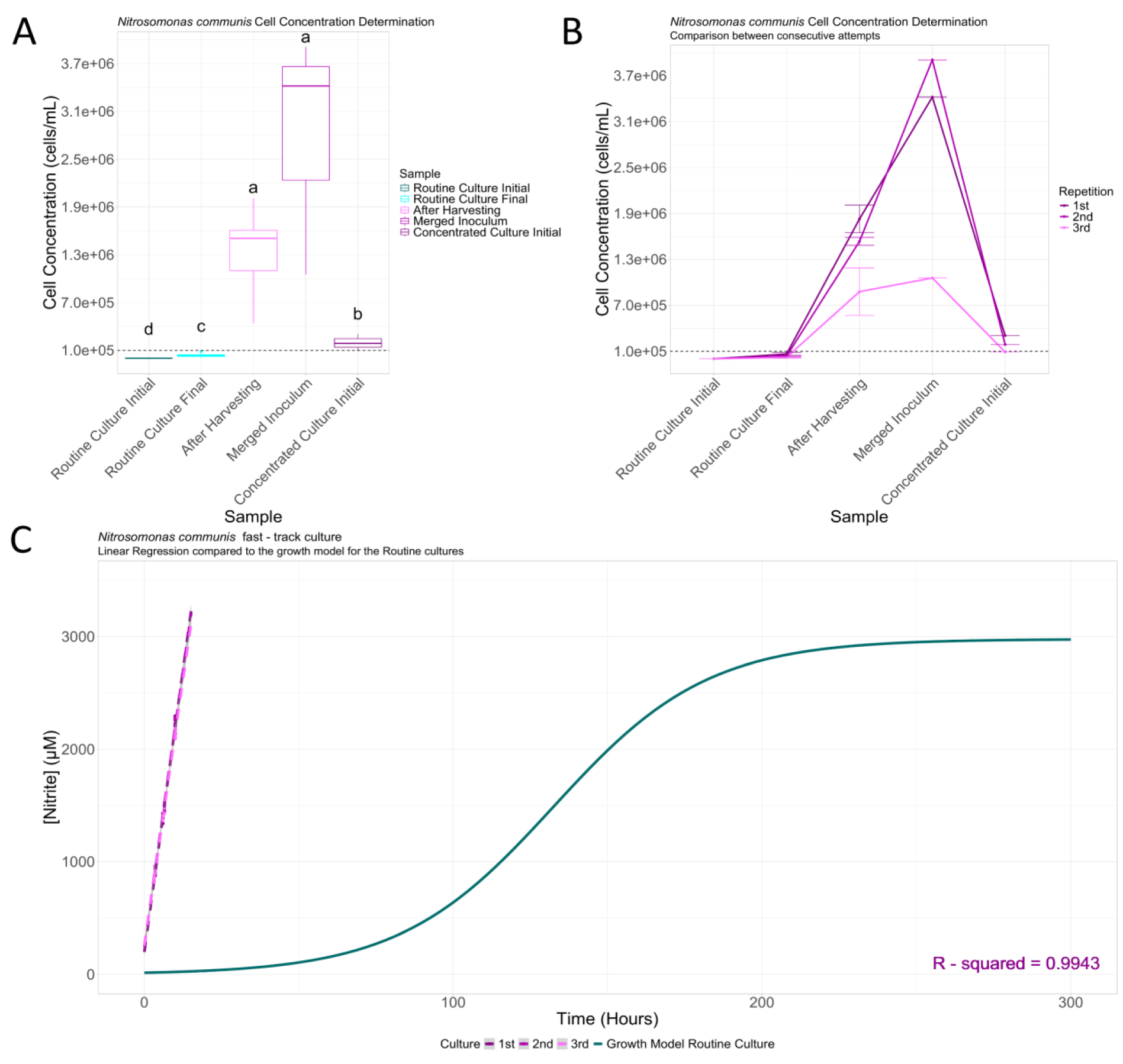
Nitrosomonas communis fast – track assay protocol standardisation. (A) Box plot of pooled qPCR results for the different points of the cell harvesting procedure. The y – axis values correspond to cell concentration, while the five different points of the procedure are plotted on the x – axis. The black dashed line represents the limit below which the box plot is not to scale. Lowercase letters above the boxes indicate the grouping according to the Kruskal – Wallis test. (B) Line plots of qPCR results between consecutive tests of the cell harvesting procedure. Different tests are represented by different colours according to the legend. The coloured error bars represent the range between -1 and +1 standard deviation from the mean value. The black dashed line indicates the limit below which the line plot is not to scale. (C) Activity curve of N. communis fast – track culture (magenta lines) compared to the activity model curve of the respective routine cultures (blue line). The points represent individual nitrite values from the biological replicates at each time-point of each fast – track culture. The R^2^ value refers to the linear regression of the pooled fast – track culture data.

**Figure S4.**
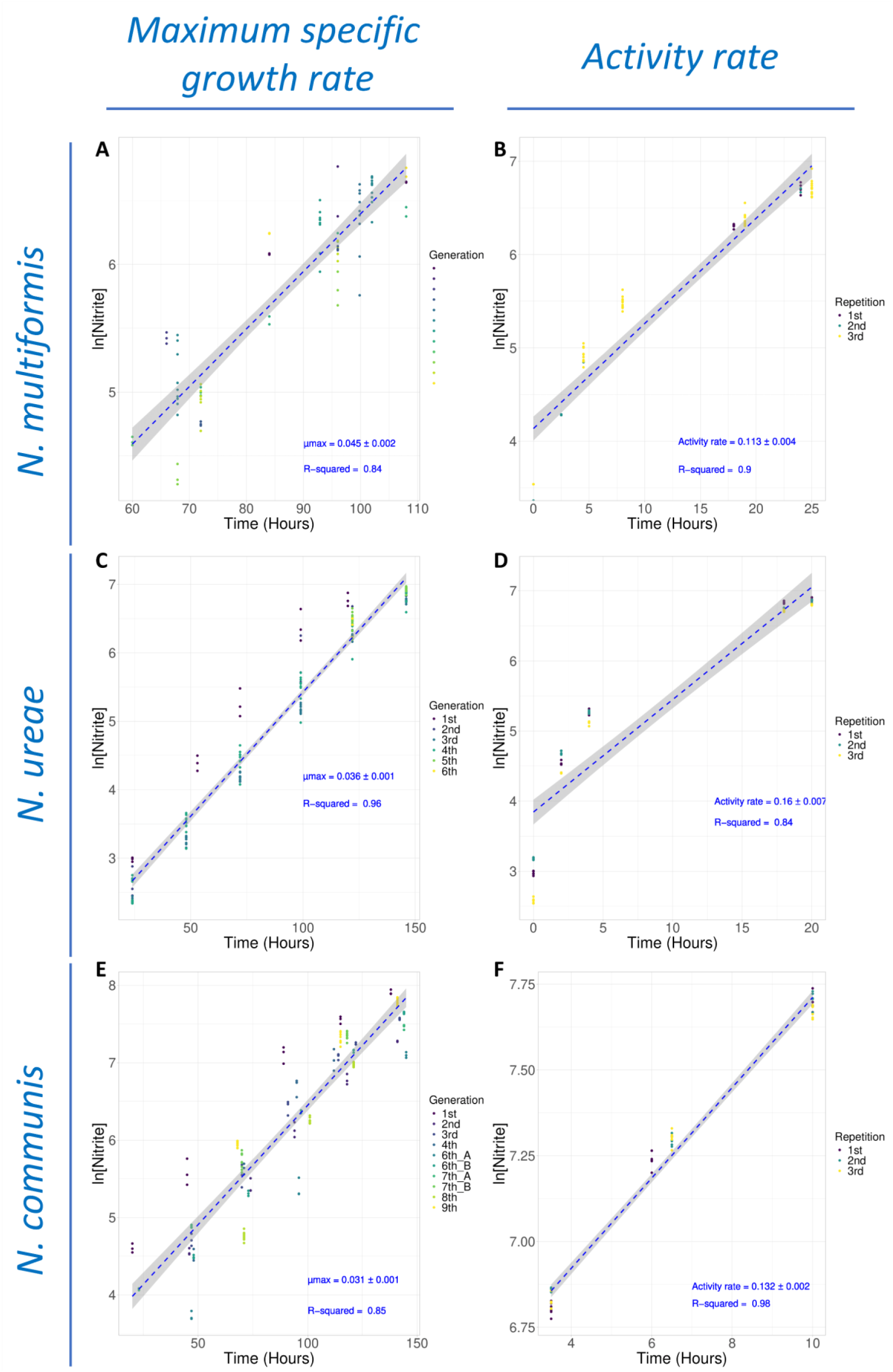
Calculation of AOB maximum specific growth and activity rates. Estimation of (A, C,E) the maximum specific growth rate (μ_max_) of AOB routine cultures and (B,D,F) the activity rate of the AOB concentrated cultures for N. multiformis (A-B), N. ureae (C-D), and N. communis (E-F). Log-transformed (ln) nitrite production values are plotted against time for consecutive generations of the AOB routine cultures or consecutive repetitions of the fast – track assay. The dotted blue line represents the linear regression curve, with shaded areas indicating the standard error. Data points for each generation or repetition are denoted by different colour, as indicated in the legend. μ_max_ and activity rate values with the respective standard errors are presented on the right side of the graphs, accompanied by the adjusted R^2^ of the respective linear regression.

**Figure S5.**
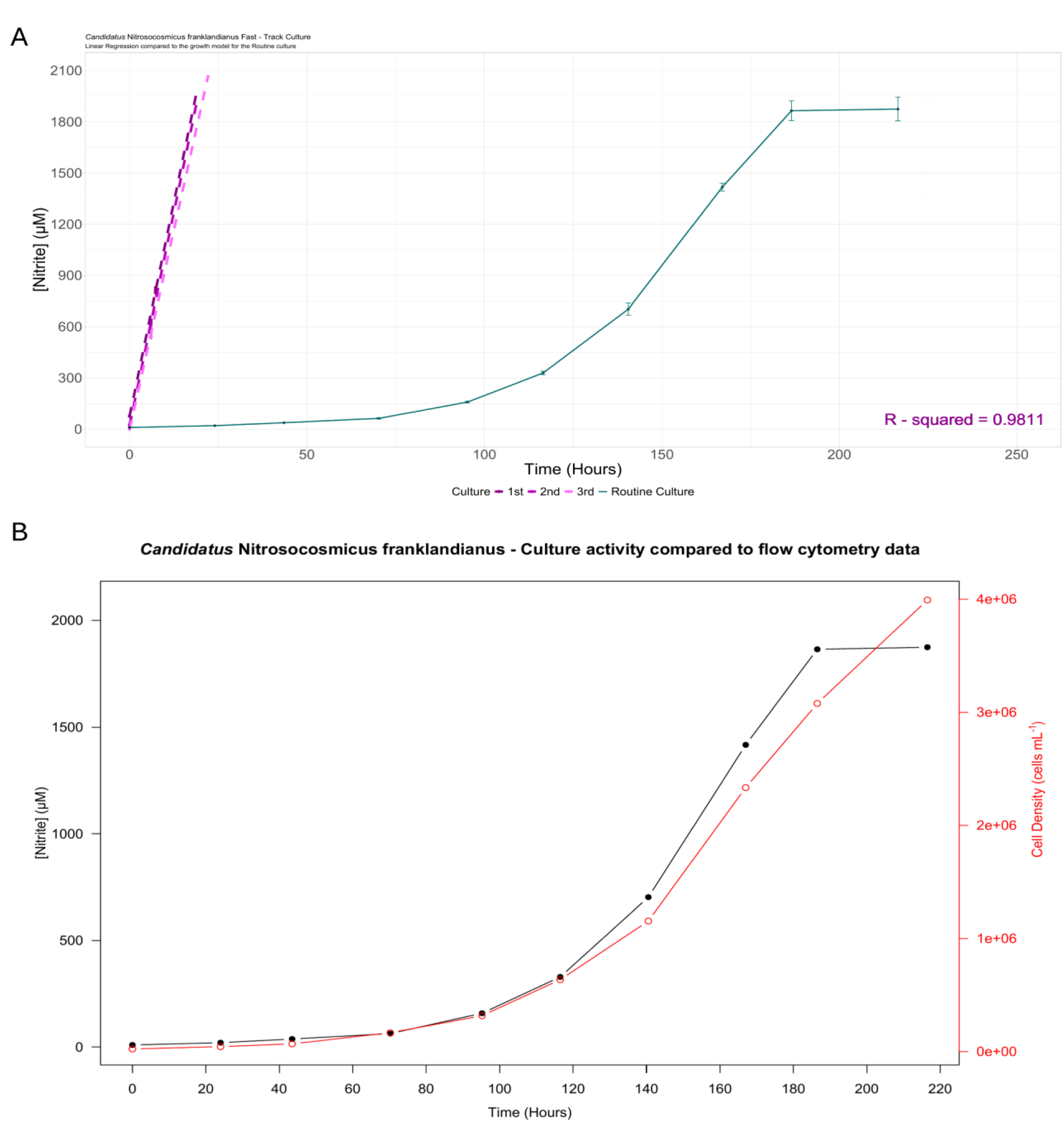
“Ca. Nitrosocosmicus franklandianus” fast – track culture standardisation. (A) Activity curves of “Ca. N. franklandianus” fast – track cultures (magenta lines) are compared to the activity model curve of the respective routine cultures (blue line). The points represent the individual nitrite values from biological replicates at each time-point of each fast – track culture. The R^2^ value refers to the linear regression of the pooled fast – track cultures data. (B) Comparison of the cell activity curve and the cell growth curve derived from flow cytometry. Points represent mean values and each y – axis, coloured differently, corresponds to the respective curve.

**Figure S6.**
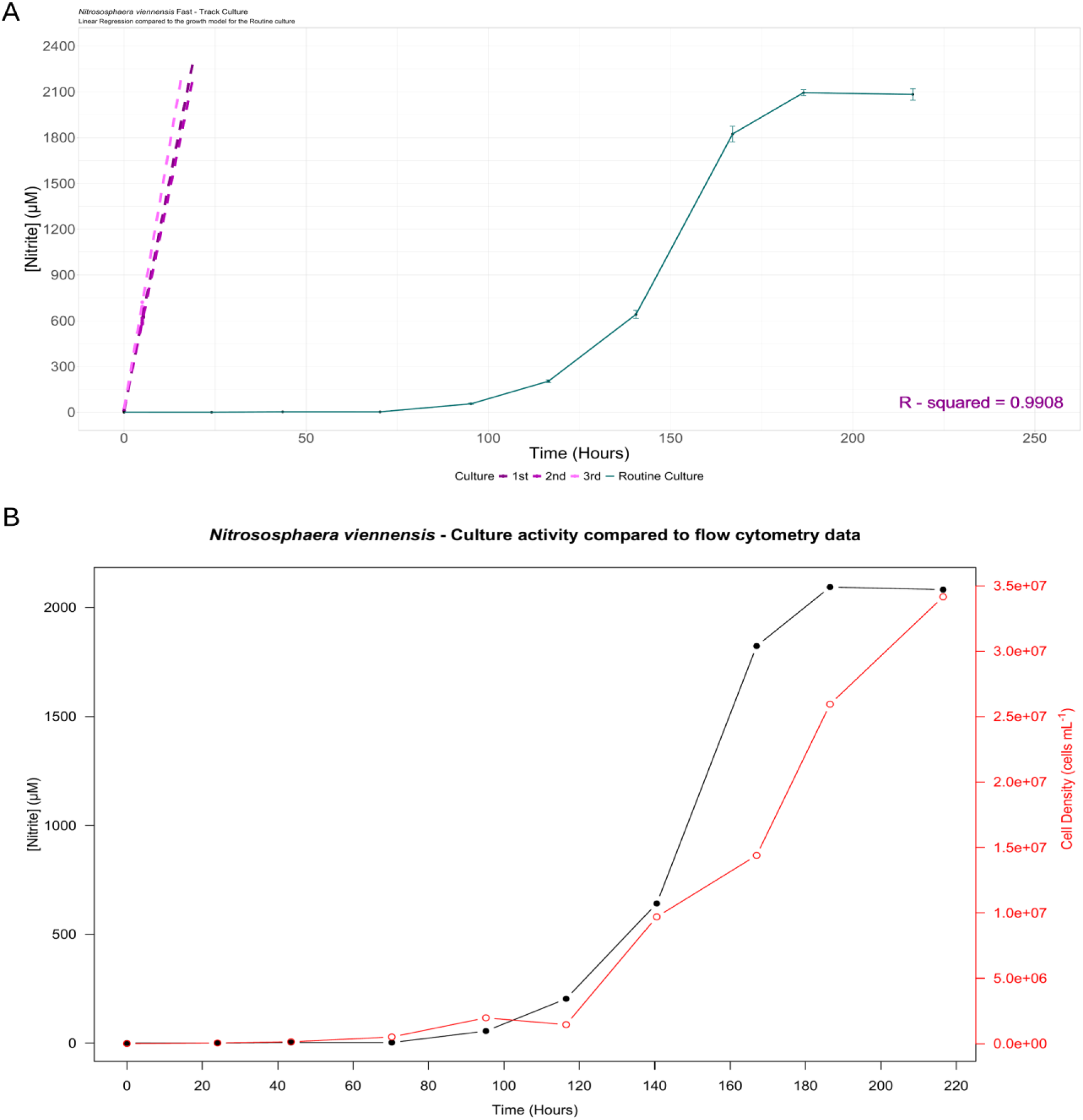
Nitrososphaera viennensis fast – track culture standardisation. (A) Activity curves of Nitrososphaera viennensis fast – track cultures (magenta lines) are compared to the activity model curve of the respective routine cultures (blue line). The points represent the individual nitrite values from biological replicates at each time-point of each fast – track culture. The R^2^ value refers to the linear regression of the pooled fast – track cultures data. (B) Comparison of the cell activity curve and the cell growth curve derived from flow cytometry. Points represent mean values and each y – axis, coloured differently, corresponds to the respective curve.

**Figure S7.**
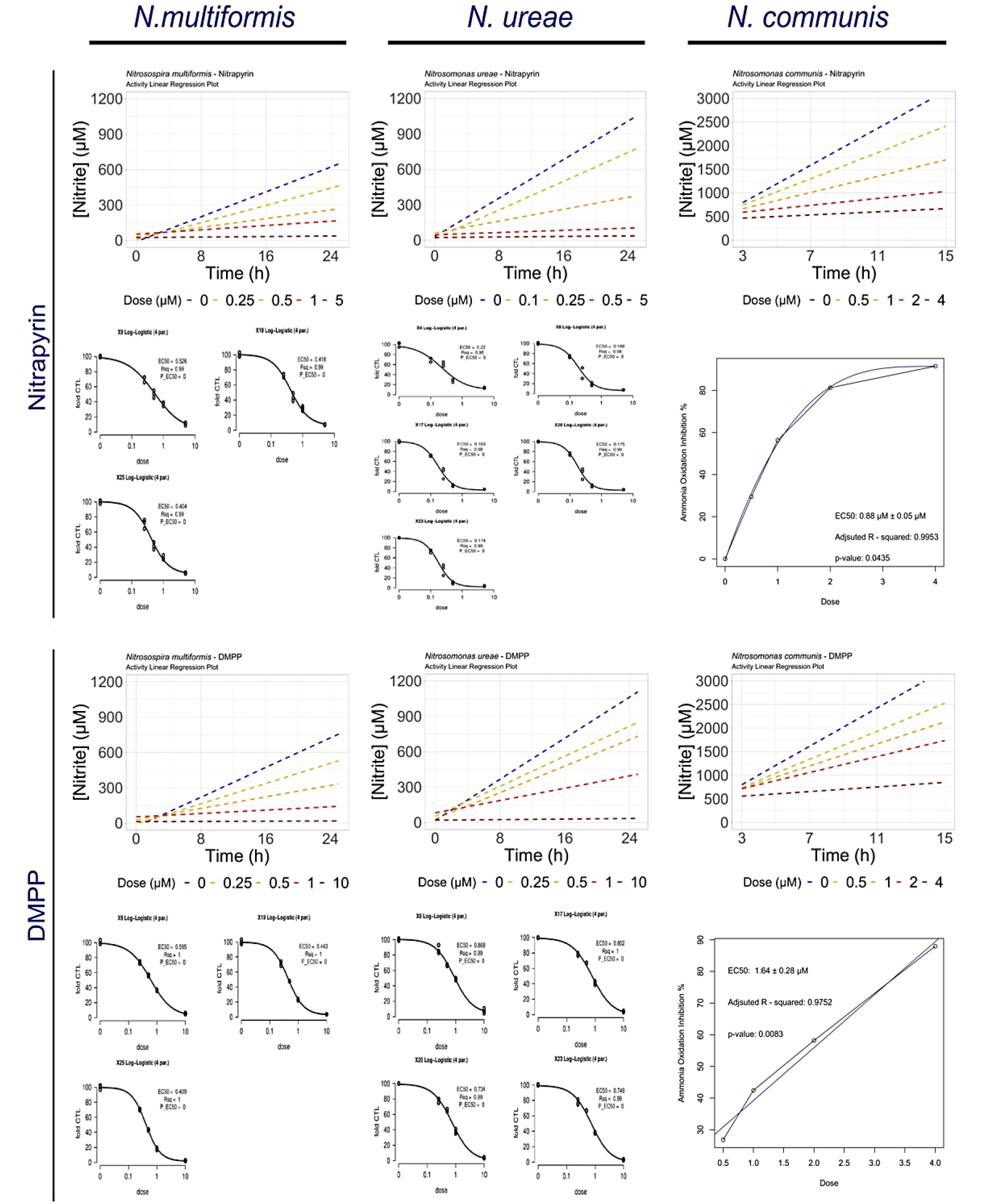
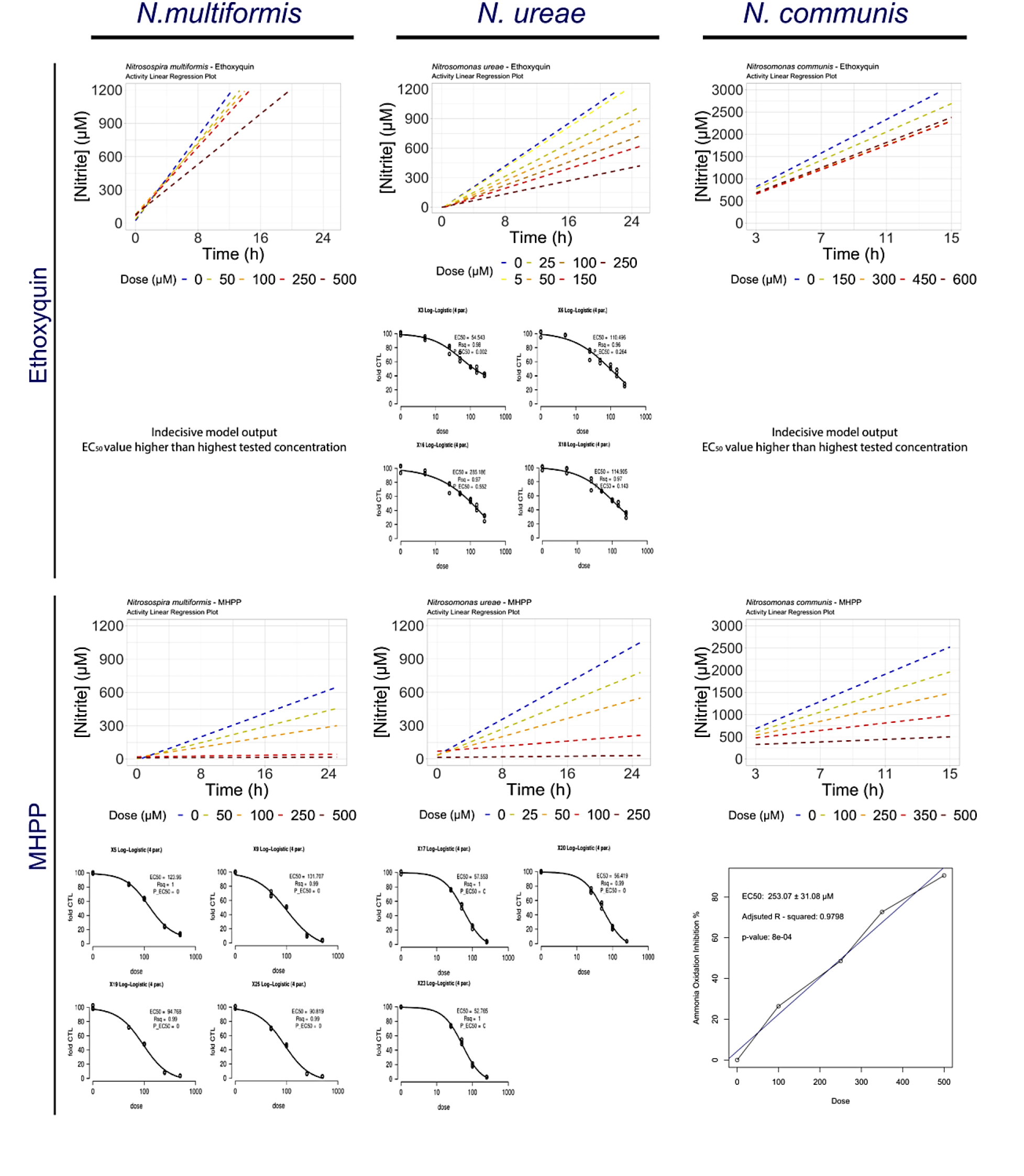
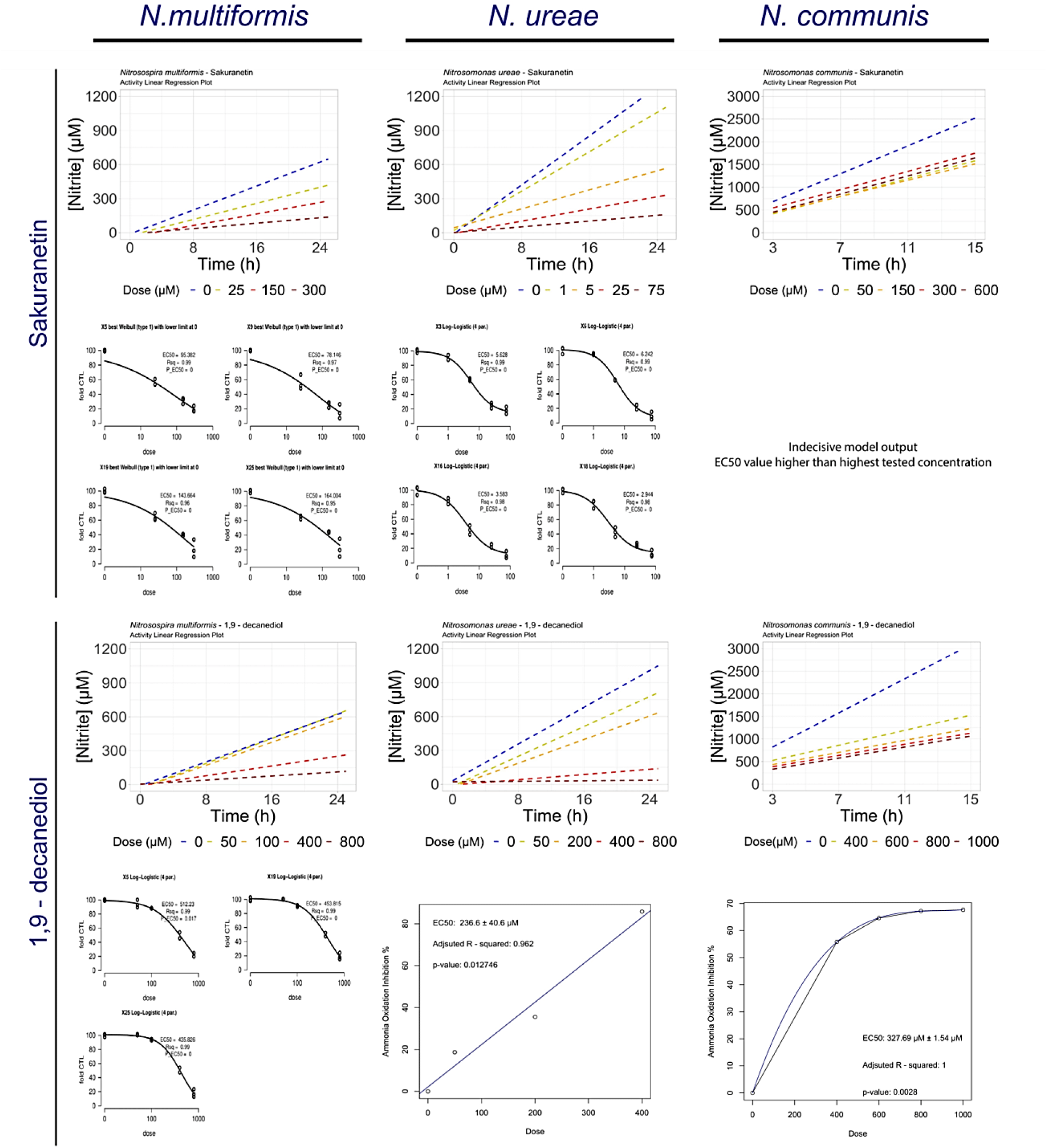
Validation of the AOB fast – track assay with pure SNIs and BNIs. The responses of the AOB strains to the tested Biological Nitrification Inhibitors (BNIs) and Synthetic Nitrification Inhibitors (SNIs) are presented. Each of the six panels of the figure corresponds to one nitrification inhibitor and the figure is divided into three columns, each representing one AOB strain. For each strain and NI combination, the ammonia oxidation activity is illustrated through a linear regression plot, comparing the AOB response to different dose rates, alongside the respective control treatment. The dashed lines in each plot represent the linear regression of nitrite values (data points not shown) at each time point for each treatment. Below the plot, a summary of the modeling of the AOB response, that led to the estimation of each EC_50_ value, is provided. For the log – logistic model, one model fit with an estimate of the EC_50_ is given for each time point. For polynomial or linear models, an aggregate model fit is presented with a final estimate of the EC_50_. Detailed information about the modeling procedure can be found in the Materials and Methods section.

**Figure S8.**
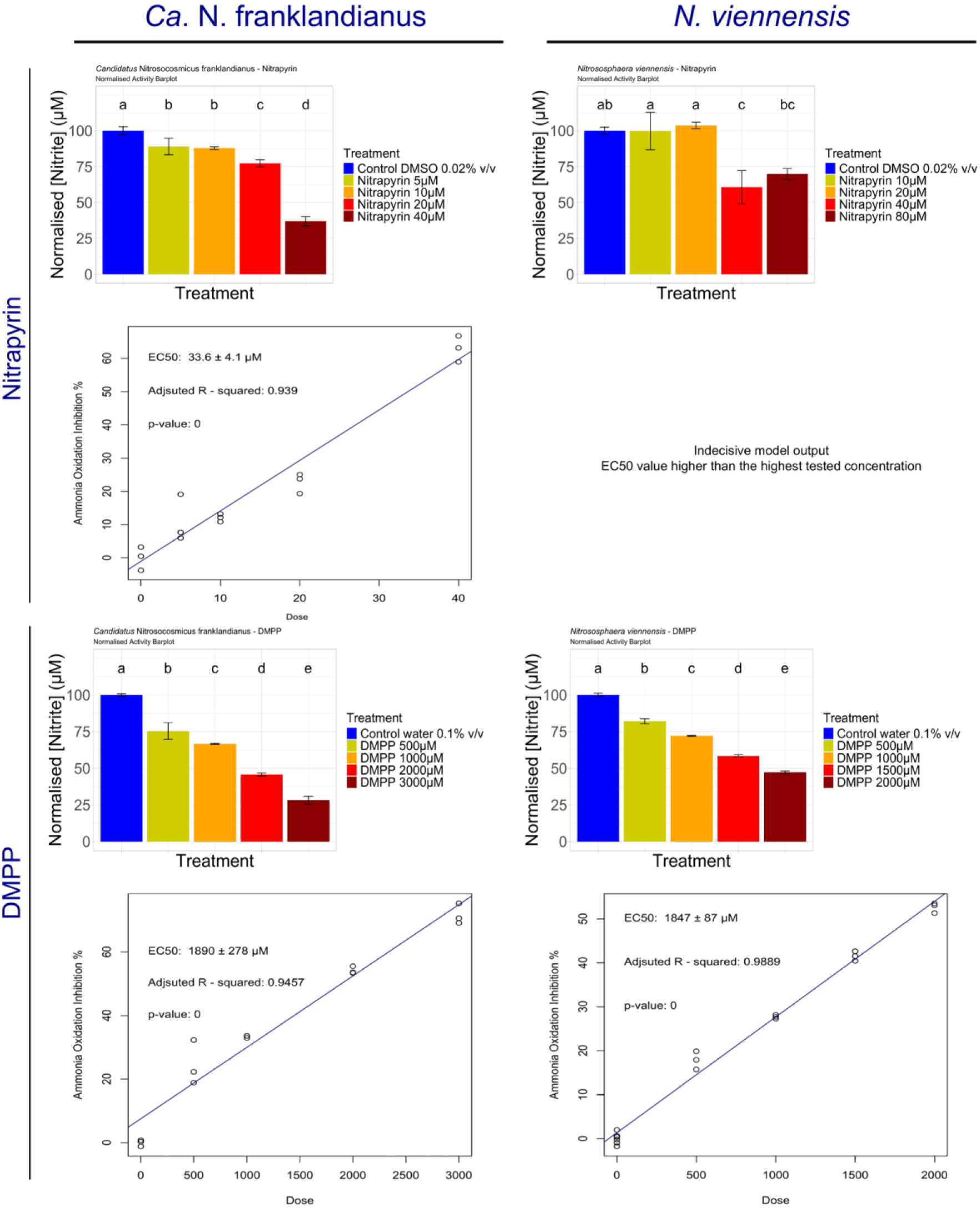
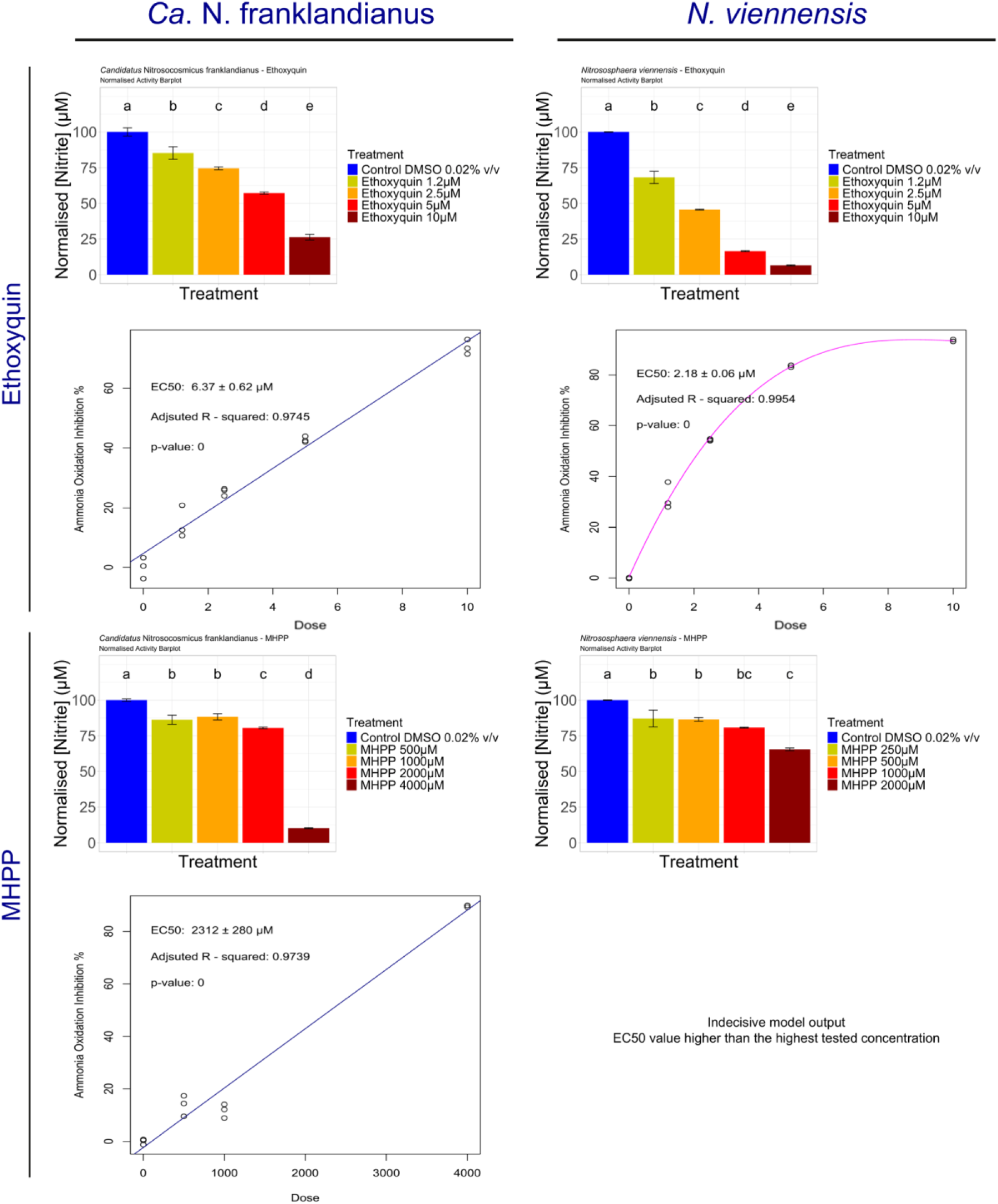
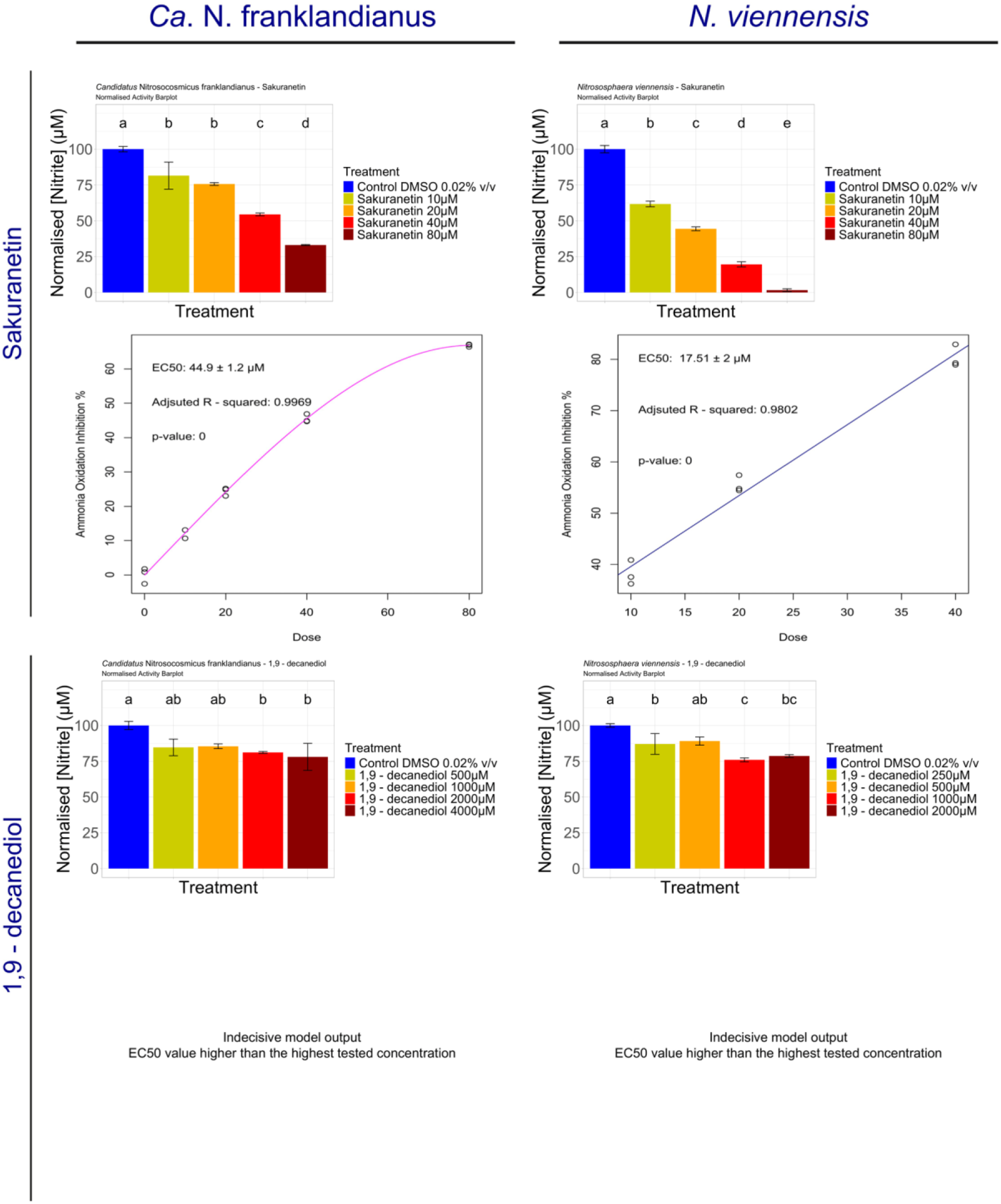
Validation of the AOA fast – track assay with pure SNIs and BNIs. The response of the AOA strains to the tested BNIs and SNIs is presented. Each of the six panels of the figure corresponds to one nitrification inhibitor and the figure is divided into two columns, each representing one AOΑ strain. For each strain and NI combination, the normalized ammonia oxidation activity data are presented in a bar plot, comparing the AOA response to different dose rates, alongside the respective control treatment. The colours of the bars correspond to different dose rates, following the legend on the right, and lowercase letters indicate the grouping of the treatments according to the Kruskal – Wallis test. Below the plot, a summary of the modeling of the AOB response, that led to the estimation of each EC_50_ value, is provided, utilizing a single model fit to estimate the EC_50_. Detailed information about the modeling procedure can be found in the Materials and Methods section.

**Figure S9.**
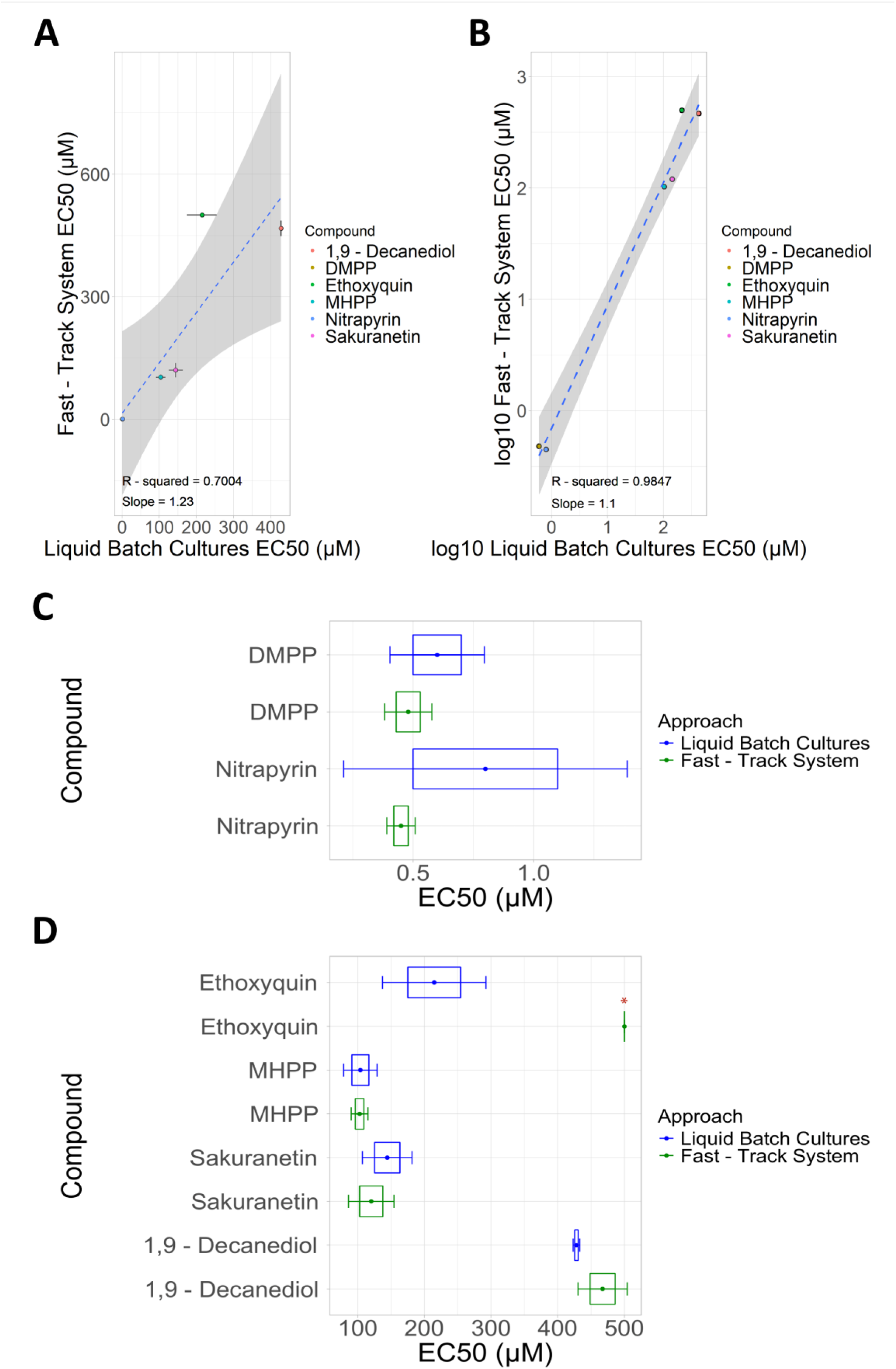
Correlation of EC_50_ values between the fast-track and established liquid batch culture systems for N. multiformis. (A) Correlation plots of EC_50_ values for the tested nitrification inhibitors and (B) transformed plot based on the log10 of the EC_50_ values. The dashed line represents the linear regression of the fitted data, and shaded areas indicate the standard error. Error bars illustrate the –1SE < EC_50_ < +1SE range. (C-D) Comparison of EC_50_ values between the fast-track and established liquid batch culture systems for the tested SNIs and BNIs. Coloured points in the middle indicate the mean EC_50_ values, while blue and green boxes show the range – 1SE < EC_50_ < +1SE. Error bars represent the range –1.96SE < EC_50_ < +1.96SE, providing a 95% confidence interval for comparison between the means. [SE: Standard Error]. Indefinite mean EC_50_ values are indicated with a red asterisk over the bar.

**Figure S10.**
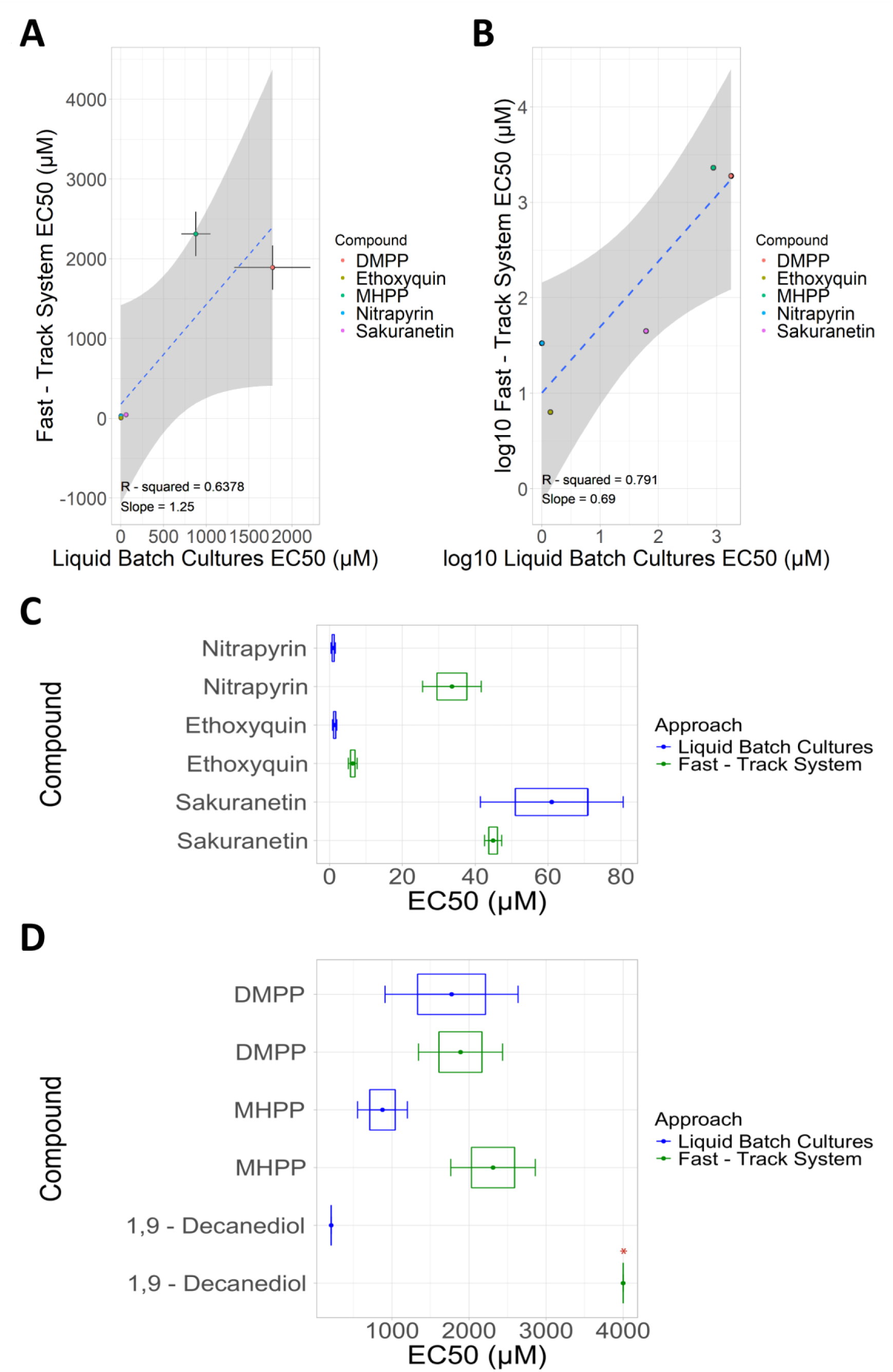
Correlation of EC_50_ values between the fast-track and established liquid batch culture systems for “Ca. N. franklandianus”. (A) Correlation plots of EC_50_ values for the tested nitrification inhibitors and (B) transformed plot based on the log10 of the EC_50_ values. The dashed line represents the linear regression of the fitted data, and shaded areas indicate the standard error. Error bars illustrate the –1SE < EC_50_ < +1SE range. (C-D) Comparison of EC_50_ values between the fast-track and established liquid batch culture systems for the tested SNIs and BNIs. Coloured points in the middle indicate the mean EC_50_ values, while blue and green boxes show the range –1SE < EC_50_ < +1SE. Error bars represent the range –1.96SE < EC_50_ < +1.96SE, providing a 95% confidence interval for comparison between the means. [SE: Standard Error]. Indefinite mean EC_50_ values are indicated with a red asterisk over the bar. 1,9 – decanediol was excluded from the analysis, as inclusion of the respective data resulted in a low-quality regression.

**Figure S11.**
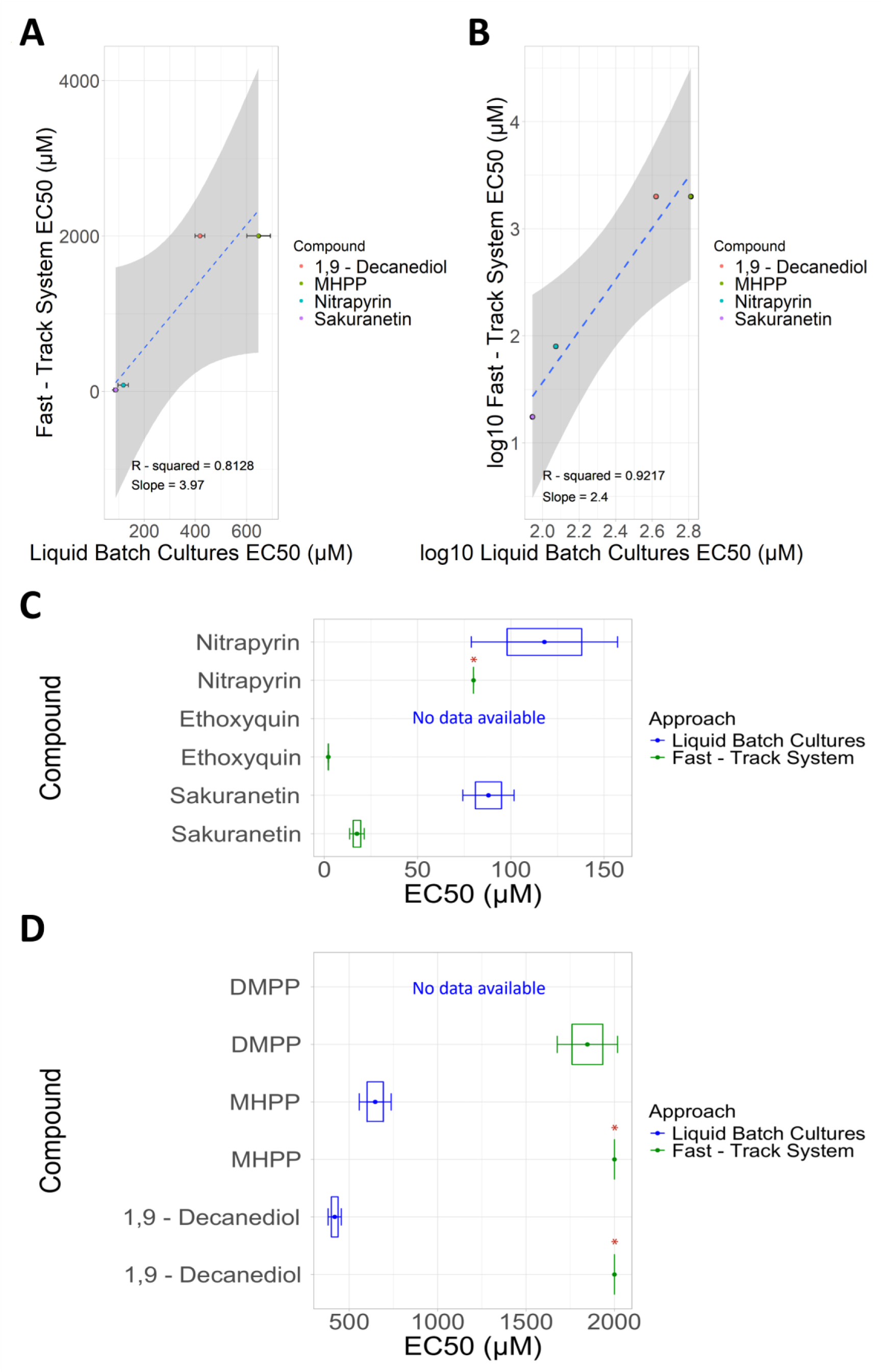
Correlation of EC_50_ values between the fast-track and established liquid batch culture systems for N. viennensis. (A) Correlation plots of EC_50_ values for the tested nitrification inhibitors and (B) transformed plot based on the log10 of the EC_50_ values. The dashed line represents the linear regression of the fitted data, and shaded areas indicate the standard error. Error bars illustrate the –1SE < EC_50_ < +1SE range. (C-D) Comparison of EC_50_ values between the fast-track and established liquid batch culture systems for the tested SNIs and BNIs. Coloured points in the middle indicate the mean EC_50_ values, while blue and green boxes show the range – 1SE < EC_50_ < +1SE. Error bars represent the range –1.96SE < EC_50_ < +1.96SE, providing a 95% confidence interval for comparison between the means. [SE: Standard Error]. Indefinite mean EC_50_ values are indicated with a red asterisk over the bar. DMPP and ethoxyquin were excluded from the analysis, as no inhibition thresholds were found in literature for this strain.

**Figure S12.**
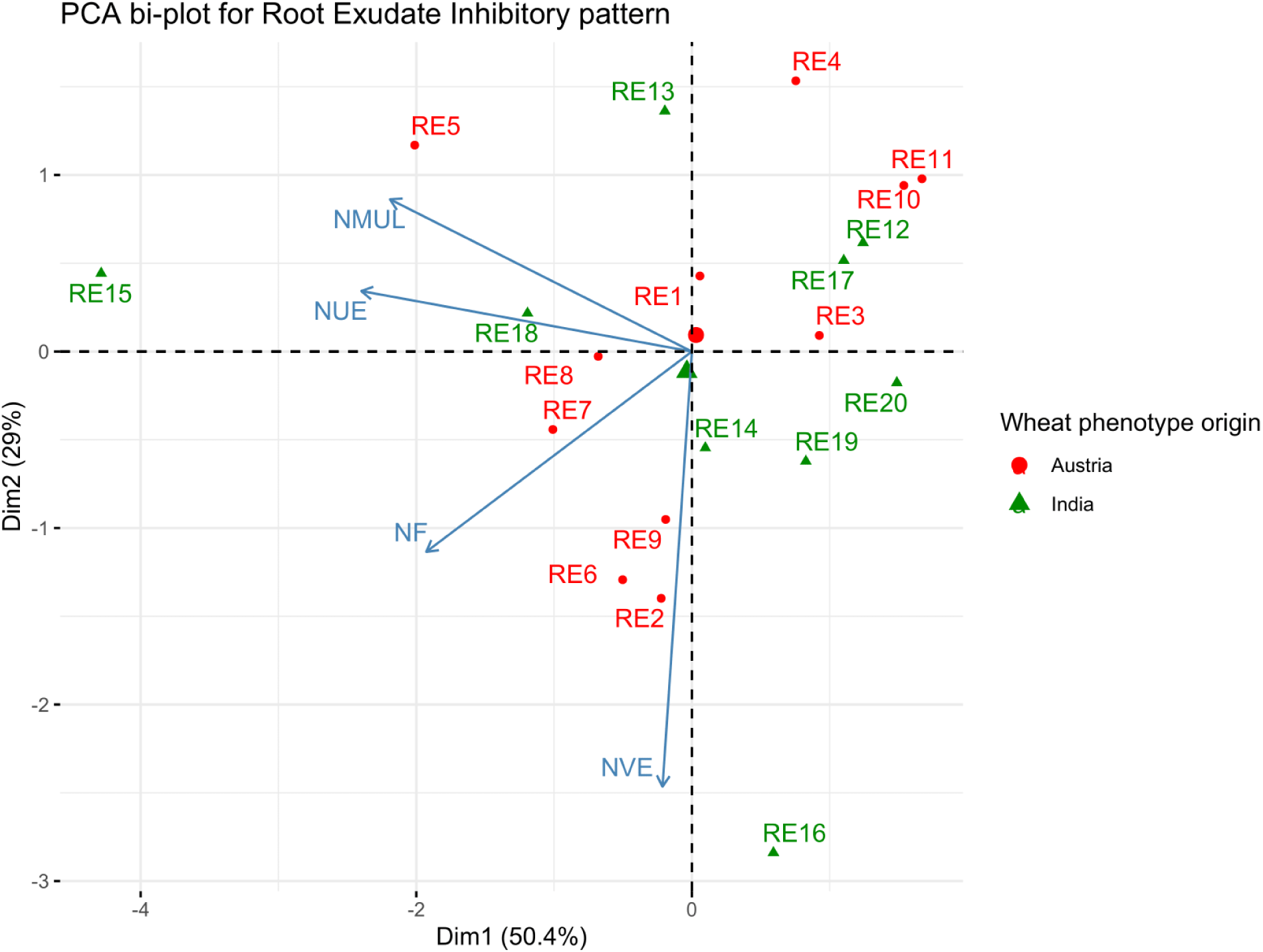
Inhibitory activity patterns of root exudates on the tested ammonia-oxidizing microorganisms (AOM). Two-dimensional representation of the root exudate inhibitory activity, derived from a four-variable dataset of the ammonia oxidation inhibition percentage values (AOI%) of each root exudate (RE) across four reporter AOM strains. Data points are color-coded based on the geographical origin of the respective genotypes. The percentage values on the axes indicate the proportion of the initial variance explained by the two principal components. The length of the light blue arrows reflects the contribution of each variable (AOM strain inhibition) to the first two dimensions, while the direction of the arrows serves as a proxy for correlation; arrows and data points on the same side indicate positively correlated variable-individual pairs. Due to limited material availability, the two most sensitive AOB (NMUL – N. multiformis, NUE – N. ureae) were included in the screening of wheat REs, along with the two AOA strains (NVE – N. viennensis, NF – “Ca. N. franklandianus”).

## References

1. Kuypers MMM, Marchant HK, Kartal B. The microbial nitrogen-cycling network. Nat Rev Microbiol. 2018; 16(5):263–76.

2. Kerou M, Alves R, Schleper C. Nitrososphaeria. In: Bergey’s Manual of systematics of Archaea and Bacteria. In 2016. p. 1–8.

3. Lehtovirta-Morley LE. Ammonia oxidation: Ecology, physiology, biochemistry and why they must all come together. FEMS Microbiol Lett. 2018; 365(9): fny058.

4. Stein LY. Insights into the physiology of ammonia-oxidizing microorganisms. Curr Opin Chem Biol. 2019; 49:9–15.

5. van Kessel MAHJ, Speth DR, Albertsen M, Nielsen PH, Op den Camp HJM, Kartal B, et al. Complete nitrification by a single microorganism. Nature. 2015; 528(7583):555–9.

6. Daims H, Lebedeva EV, Pjevac P, Han P, Herbold C, Albertsen M, et al. Complete nitrification by *Nitrospira* bacteria. Nature. 2015; 528(7583):504–9.

7. Erisman JW, Sutton MA, Galloway J, Klimont Z, Winiwarter W. How a century of ammonia synthesis changed the world. Nat Geosci. 2008;1(10):636–9.

8. Stein LY. Agritech to Tame the Nitrogen Cycle. Cold Spring Harb Perspect Biol. 2024;16(3): a041668.

9. Battye W, Aneja VP, Schlesinger WH. Is nitrogen the next carbon? Earths Future. 2017; 5(9):894–904.

10. Thomson AJ, Giannopoulos G, Pretty J, Baggs EM, Richardson DJ. Biological sources and sinks of nitrous oxide and strategies to mitigate emissions. Philos Trans R Soc B Biol Sci. 2012; 367(1593):1157–68.

11. Wang X, Bai J, Xie T, Wang W, Zhang G, Yin S, et al. Effects of biological nitrification inhibitors on nitrogen use efficiency and greenhouse gas emissions in agricultural soils: A review. Ecotoxicol Environ Saf. 2021; 220:112338.

12. Huang L, Chakrabarti S, Cooper J, Perez A, John SM, Daroub SH, et al. Ammonia-oxidizing archaea are integral to nitrogen cycling in a highly fertile agricultural soil. ISME Commun. 2021;1(1):19.

13. Lin L, St Clair S, Gamble GD, Crowther CA, Dixon L, Bloomfield FH, et al. Nitrate contamination in drinking water and adverse reproductive and birth outcomes: a systematic review and meta-analysis. Sci Rep. 2023;13(1):563.

14. Prosser JI, Hink L, Gubry-Rangin C, Nicol GW. Nitrous oxide production by ammonia oxidizers: Physiological diversity, niche differentiation and potential mitigation strategies. Glob Change Biol. 2020; 26(1):103–18.

15. Stein LY. The long-term relationship between microbial metabolism and greenhouse gases. Trends Microbiol. 2020; 28(6):500–11.

16. van Groenigen JW, Huygens D, Boeckx P, Kuyper TW, Lubbers IM, Rütting T, et al. The soil N cycle: New insights and key challenges. Soil. 11;1(1):235–56.

17. Kozlowski JA, Stieglmeier M, Schleper C, Klotz MG, Stein LY. Pathways and key intermediates required for obligate aerobic ammonia-dependent chemolithotrophy in bacteria and Thaumarchaeota. ISME J. 2016;10(8):1836–45.

18. Prasad R, Power JF. Nitrification inhibitors for agriculture, health, and the environment. In: Advances in Agronomy. Elsevier. 1995; 233–81.

19. Subbarao GV, Ito O, Sahrawat KL, Berry WL, Nakahara K, Ishikawa T, et al. Scope and strategies for Regulation of Nitrification in Agricultural Systems—Challenges and Opportunities. Crit Rev Plant Sci. 2006;25(4):303–35.

20. Abalos D, Jeffery S, Sanz-Cobena A, Guardia G, Vallejo A. Meta-analysis of the effect of urease and nitrification inhibitors on crop productivity and nitrogen use efficiency. Agric Ecosyst Environ. 2014; 189:136–44.

21. Ruser R, Schulz R. The effect of nitrification inhibitors on the nitrous oxide (N_2_O) release from agricultural soils—a review. J Plant Nutr Soil Sci. 2015;178(2):171–88.

22. Papadopoulou ES, Tsachidou B, Sułowicz S, Menkissoglu-Spiroudi U, Karpouzas DG. Land spreading of wastewaters from the fruit-packaging industry and potential effects on soil microbes: effects of the antioxidant ethoxyquin and its metabolites on ammonia oxidizers. Appl Environ Microbiol. 2016;82(2):747–55.

23. Taggert BI, Walker C, Chen D, Wille U. Substituted 1,2,3-triazoles: A new class of nitrification inhibitors. Sci Rep. 2021;11(1):14980.

24. Beeckman F, Drozdzecki A, De Knijf A, Corrochano-Monsalve M, Bodé S, Blom P, et al. Drug discovery-based approach identifies new nitrification inhibitors. J Environ Manage. 2023; 346:118996.

25. Nardi P, Laanbroek HJ, Nicol GW, Renella G, Cardinale M, Pietramellara G, et al. Biological nitrification inhibition in the rhizosphere: determining interactions and impact on microbially mediated processes and potential applications. FEMS Microbiol Rev. 2020; 44(6):874–908.

26. Bachtsevani E, Papazlatani CV, Rousidou C, Lampronikou E, Menkissoglu-Spiroudi U, Nicol GW, et al. Effects of the nitrification inhibitor 3,4-dimethylpyrazole phosphate (DMPP) on the activity and diversity of the soil microbial community under contrasting soil pH. Biol Fertil Soils. 2021;57(8):1117–35.

27. Coskun D, Britto DT, Shi W, Kronzucker HJ. Nitrogen transformations in modern agriculture and the role of biological nitrification inhibition. Nat Plants. 2017;3(6):1–10.

28. Ghatak A, Chaturvedi P, Waldherr S, Subbarao GV, Weckwerth W. PANOMICS at the interface of root–soil microbiome and BNI. Trends Plant Sci. 2023;28(1):106–22.

29. Subbarao GV, Nakahara K, Ishikawa T, Ono H, Yoshida M, Yoshihashi T, et al. Biological nitrification inhibition (BNI) activity in sorghum and its characterization. Plant Soil. 2013;366(1):243–59.

30. Zakir HAKM, Subbarao GV, Pearse SJ, Gopalakrishnan S, Ito O, Ishikawa T, et al. Detection, isolation and characterization of a root-exuded compound, methyl 3-(4-hydroxyphenyl) propionate, responsible for biological nitrification inhibition by sorghum (Sorghum bicolor). New Phytol. 2008;180(2):442–51.

31. Sun L, Lu Y, Yu F, Kronzucker HJ, Shi W. Biological nitrification inhibition by rice root exudates and its relationship with nitrogen-use efficiency. New Phytol. 2016;212(3):646–56.

32. Kaur-Bhambra J, Wardak DLR, Prosser JI, Gubry-Rangin C. Revisiting plant biological nitrification inhibition efficiency using multiple archaeal and bacterial ammonia-oxidising cultures. Biol Fertil Soils. 2022;58(3):241–9.

33. Kolovou M, Panagiotou D, Süße L, Loiseleur O, Williams S, Karpouzas DG, et al. Assessing the activity of different plant-derived molecules and potential biological nitrification inhibitors on a range of soil ammonia- and nitrite-oxidizing strains. Appl Environ Microbiol. 2023; 89(11): e01380–23.

34. Subbarao GV, Nakahara K, Hurtado MP, Ono H, Moreta DE, Salcedo AF, et al. Evidence for biological nitrification inhibition in Brachiaria pastures. Proc Natl Acad Sci. 2009;106(41):17302–7.

35. Villegas D, Arevalo A, Nuñez J, Mazabel J, Subbarao G, Rao I, et al. Biological Nitrification Inhibition (BNI): Phenotyping of a core germplasm collection of the tropical forage grass *Megathyrsus maximus* under greenhouse conditions. Front Plant Sci. 2020;11: 820.

36. Shen T, Stieglmeier M, Dai J, Urich T, Schleper C. Responses of the terrestrial ammonia-oxidizing archaeon *Ca.* Nitrososphaera viennensis and the ammonia-oxidizing bacterium *Nitrosospira multiformis* to nitrification inhibitors. FEMS Microbiol Lett. 2013; 344(2):121–9.

37. Papadopoulou ES, Bachtsevani E, Lampronikou E, Adamou E, Katsaouni A, Vasileiadis S, et al. Comparison of novel and established nitrification inhibitors relevant to agriculture on soil ammonia- and nitrite-oxidizing isolates. Front Microbiol. 2020; 11:581283.

38. Subbarao GV, Ishikawa T, Ito O, Nakahara K, Wang HY, Berry WL. A bioluminescence assay to detect nitrification inhibitors released from plant roots: a case study with Brachiaria humidicola. Plant Soil. 2006; 288(1):101–12.

39. Subbarao GV, Tomohiro B, Masahiro K, Osamu I, Samejima H, Wang HY, et al. Can biological nitrification inhibition (BNI) genes from perennial *Leymus racemosus* (Triticeae) combat nitrification in wheat farming? Plant Soil. 2007; 299(1):55–64.

40. Iizumi T, Mizumoto M, Nakamura K. A bioluminescence assay using *Nitrosomonas europaea* for rapid and sensitive detection of nitrification inhibitors. Appl Environ Microbiol. 1998; 64(10):3656–62.

41. O’Sullivan CA, Fillery IRP, Roper MM, Richards RA. Identification of several wheat landraces with biological nitrification inhibition capacity. Plant Soil. 2016; 404(1/2):61–74.

42. Schleper C, Nicol GW. Ammonia-Oxidising Archaea – Physiology, Ecology and Evolution. In: Poole RK, editor. Advances in Microbial Physiology. Academic Press. 2010;1-41.

43. Skinner FA, Walker N. Growth of *Nitrosomonas europaea* in batch and continuous culture. Arch Für Mikrobiol. 1961;38(4):339–49.

44. Koops HP, Böttcher B, Möller UC, Pommerening-Röser A, Stehr G. Classification of eight new species of ammonia-oxidizing bacteria: *Nitrosomonas communis* sp. nov., *Nitrosomonas ureae* sp. nov., *Nitrosomonas aestuarii* sp. nov., *Nitrosomonas marina* sp. nov., *Nitrosomonas nitrosa* sp. nov., *Nitrosomonas eutropha* sp. nov., Nitrosomonas oligotropha sp. nov. and Nitrosomonas halophila sp. nov. Microbiology. 1991;137(7):1689–99.

45. Reyes C, Hodgskiss LH, Baars O, Kerou M, Bayer B, Schleper C, et al. Copper limiting threshold in the terrestrial ammonia oxidizing archaeon *Nitrososphaera viennensis*. Res Microbiol. 2020;171(3):134–42.

46. Zerulla W, Barth T, Dressel J, Erhardt K, Horchler von Locquenghien K, Pasda G, et al. 3,4-Dimethylpyrazole phosphate (DMPP) – a new nitrification inhibitor for agriculture and horticulture. Biol Fertil Soils. 2001;34(2):79–84.

47. Goring C a. I. Control of nitrification by 2-chloro-6-(trichloro-methyl) pyridine. Soil Sci. 1962; 93(3):211.

48. Shinn MB. ACS Publications. American Chemical Society; 1941. Colorimetric method for determination of nitrate. https://pubs.acs.org/doi/abs/10.1021/i560089a010

49. Wright CL, Schatteman A, Crombie AT, Murrell JC, Lehtovirta-Morley LE. Inhibition of ammonia monooxygenase from ammonia-oxidizing archaea by linear and aromatic alkynes. Appl Environ Microbiol. 2020; 86(9): e02388–19.

50. Zhao J, Bello MO, Meng Y, Prosser JI, Gubry-Rangin C. Selective inhibition of ammonia oxidising archaea by simvastatin stimulates growth of ammonia oxidising bacteria. Soil Biol Biochem. 2020; 141:107673.

51. Ghatak A, Schindler F, Bachmann G, Engelmeier D, Bajaj P, Brenner M, et al. Root exudation of contrasting drought-stressed pearl millet genotypes conveys varying biological nitrification inhibition (BNI) activity. Biol Fertil Soils. 2022; 58(3):291–306.

52. Rotthauwe JH, Witzel KP, Liesack W. The ammonia monooxygenase structural gene amoA as a functional marker: molecular fine-scale analysis of natural ammonia-oxidizing populations. Appl Environ Microbiol. 1997; 63(12):4704–12.

53. Norton JM, Klotz MG, Stein LY, Arp DJ, Bottomley PJ, Chain PSG, et al. Complete genome sequence of *Nitrosospira multiformis*, an ammonia-oxidizing bacterium from the soil environment. Appl Environ Microbiol. 2008; 74(11):3559–72.

54. Kozlowski JA, Kits KD, Stein LY. Complete Genome Sequence of *Nitrosomonas ureae* strain Nm10, an oligotrophic group 6a Nitrosomonad. Genome Announc. 2016; 4(2):10.1128/genomea.00094-16.

55. Kozlowski JA, Kits KD, Stein LY. Genome sequence of *Nitrosomonas communis* strain Nm2, a mesophilic ammonia-oxidizing bacterium isolated from mediterranean soil. Genome Announc. 2016; 4(1):10.1128/genomea.01541-15.

56. Malits A, Ibarbalz FM, Martín J, Flombaum P. Higher biotic than abiotic natural variability of the plankton ecosystem revealed by a time series along a subantarctic transect. J Mar Syst. 2023; 238:103843.

57. Caglar MU, Teufel AI, Wilke CO. Sicegar: R package for sigmoidal and double-sigmoidal curve fitting. PeerJ. 2018; 6: e4251.

58. Powell SJ, Prosser JI. Effect of copper on inhibition by nitrapyrin of growth ofNitrosomonas europaea. Curr Microbiol. 1986;14(3):177–9.

59. Brook I. Inoculum Effect. Rev Infect Dis. 1989;11(3):361–8.

60. Bottery MJ, Pitchford JW, Friman VP. Ecology and evolution of antimicrobial resistance in bacterial communities. ISME J. 2021;15(4):939–48.

61. Vannelli T, Hooper AB. Oxidation of nitrapyrin to 6-chloropicolinic acid by the ammonia-oxidizing bacterium *Nitrosomonas europaea*. Appl Environ Microbiol. 1992; 58(7):2321–5.

62. Powell SJ, Prosser JI. The effect of nitrapyrin and chloropicolinic acid on ammonium oxidation by *Nitrosomonas europaea*. FEMS Microbiol Lett. 1985; 28(1):51–4.

63. Lin Y, Ye G, Luo J, Di HJ, Liu D, Fan J, et al. *Nitrosospira* cluster 8a plays a predominant role in the nitrification process of a subtropical ultisol under long-term inorganic and organic fertilization. Appl Environ Microbiol. 2018; 84(18): e01031–18.

64. Cassman NA, Soares JR, Pijl A, Lourenço KS, van Veen JA, Cantarella H, et al. Nitrification inhibitors effectively target N_2_O-producing *Nitrosospira* spp. in tropical soil. Environ Microbiol. 2019; 21(4):1241–54.

65. Zorz JK, Kozlowski JA, Stein LY, Strous M, Kleiner M. Comparative proteomics of three species of ammonia-oxidizing bacteria. Front Microbiol. 2018; 9:938.

66. Sedlacek CJ, McGowan B, Suwa Y, Sayavedra-Soto L, Laanbroek HJ, Stein LY, et al. A Physiological and genomic Comparison of *Nitrosomonas* cluster 6a and 7 ammonia-oxidizing bacteria. Microb Ecol. 2019 Nov 1;78(4):985–94.

67. Alves RJE, Minh BQ, Urich T, von Haeseler A, Schleper C. Unifying the global phylogeny and environmental distribution of ammonia-oxidising archaea based on amoA genes. Nat Commun. 2018; 9(1):1517.

68. Lehtovirta-Morley LE, Ross J, Hink L, Weber EB, Gubry-Rangin C, Thion C, et al. Isolation of ‘*Candidatus* Nitrosocosmicus franklandus’, a novel ureolytic soil archaeal ammonia oxidiser with tolerance to high ammonia concentration. FEMS Microbiol Ecol. 2016; 92(5): fiw057.

69. Han S, Kim S, Sedlacek CJ, Farooq A, Song C, Lee S, et al. Adaptive traits of *Nitrosocosmicus* clade ammonia-oxidizing archaea. mBio. 2024; 0(0): e02169–24.

70. Lee UJ, Gwak JH, Choi S, Jung MY, Lee TK, Ryu H, et al. “*Ca*. Nitrosocosmicus” members are the dominant archaea associated with plant rhizospheres. mSphere. 2024;0(0): e00821-24.

71. Belser LW, Schmidt EL. Inhibitory effect of nitrapyrin on three genera of ammonia-oxidizing nitrifiers. Appl Environ Microbiol. 1981;41(3):819–21.

72. Beeckman F, Drozdzecki A, De Knijf A, Audenaert D, Beeckman T, Motte H. High-throughput assays to identify archaea-targeting nitrification inhibitors. Front Plant Sci. 2024 J; 14:1283047.

73. Liu L, Liu M, Jiang Y, Lin W, Luo J. Physiological and genomic analysis of “*Candidatus* Nitrosocosmicus agrestis”, an ammonia tolerant ammonia-oxidizing archaeon from vegetable soil. bioRxiv. 2019; 2019.12.11.872556.

74. Kerou M, Offre P, Valledor L, Abby SS, Melcher M, Nagler M, et al. Proteomics and comparative genomics of *Nitrososphaera viennensis* reveal the core genome and adaptations of archaeal ammonia oxidizers. Proc Natl Acad Sci. 2016; 113(49): E7937–46.

75. Jung MY, Kim JG, Sinninghe Damsté JS, Rijpstra WIC, Madsen EL, Kim SJ, et al. A hydrophobic ammonia-oxidizing archaeon of the *Nitrosocosmicus* clade isolated from coal tar-contaminated sediment. Environ Microbiol Rep. 2016; 8(6):983–92.

76. Sauder LA, Albertsen M, Engel K, Schwarz J, Nielsen PH, Wagner M, et al. Cultivation and characterization of *Candidatus* Nitrosocosmicus exaquare, an ammonia-oxidizing archaeon from a municipal wastewater treatment system. ISME J. 2017;11(5):1142–57.

77. Alves RJE, Kerou M, Zappe A, Bittner R, Abby SS, Schmidt HA, et al. Ammonia oxidation by the arctic terrestrial Thaumarchaeote *Candidatus* Nitrosocosmicus arcticus Is stimulated by increasing temperatures. Front Microbiol. 2019; 10:1571.

78. von Kügelgen A, Cassidy CK, van Dorst S, Pagani LL, Batters C, Ford Z, et al. Membraneless channels sieve cations in ammonia-oxidizing marine archaea. Nature. 2024; 630(8015):230–6.

79. Ma W, Tang S, Dengzeng Z, Zhang D, Zhang T, Ma X. Root exudates contribute to belowground ecosystem hotspots: A review. Front Microbiol. 2022; 13:937940.

80. Petroli CD, Subbarao GV, Burgueño JA, Yoshihashi T, Li H, Duran JF, et al. Genetic variation among elite inbred lines suggests potential to breed for BNI-capacity in maize. Sci Rep. 2023; 13:13422.

81. Yahya M, Islam E ul, Rasul M, Farooq I, Mahreen N, Tawab A, et al. Differential root exudation and architecture for improved growth of wheat mediated by phosphate solubilizing bacteria. Front Microbiol. 2021; 12:744094.

82. Canarini A, Kaiser C, Merchant A, Richter A, Wanek W. Root exudation of primary metabolites: Μechanisms and their roles in plant responses to environmental stimuli. Front Plant Sci. 2019;10: 157.

83. Upadhyay SK, Srivastava AK, Rajput VD, Chauhan PK, Bhojiya AA, Jain D, et al. Root exudates: Mechanistic insight of plant growth promoting rhizobacteria for sustainable crop production. Front Microbiol. 2022;13: :916488.

## References

Caglar, M. U., Teufel, A. I., & Wilke, C. O. (2018). Sicegar: R package for sigmoidal and double-sigmoidal curve fitting. PeerJ, 6, e4251. 10.7717/peerj.4251

de Mendiburu F (2023). _agricolae: Statistical Procedures for Agricultural Research_. R package version 1.3–7, https://CRAN.R-project.org/package=agricolae

Galili T. (2015). dendextend: an R package for visualizing, adjusting and comparing trees of hierarchical clustering. Bioinformatics (Oxford, England), 31(22), 3718–3720. 10.1093/bioinformatics/btv428

Greenwell, B. M., Kabban, C. M. S. (2014). “investr: An R Package for Inverse Estimation.” The R Journal, 6(1), 90–100. 10.32614/RJ-2014-009

Gu, Z., Eils, R., & Schlesner, M. (2016). Complex heatmaps reveal patterns and correlations in multidimensional genomic data. Bioinformatics (Oxford, England), 32(18), 2847–2849. 10.1093/bioinformatics/btw313

Kassambara, A., Mundt, F. (2020). _factoextra: Extract and Visualize the Results of Multivariate Data Analyses_. R package version 1.0.7. https://CRAN.R-project.org/package=factoextra

Kassambara, A. (2023a). ggpubr: ’ggplot2’ Based Publication Ready Plots_. R package version 0.6.0, https://CRAN.R-project.org/package=ggpubr

Kassambara, A. (2023b). rstatix: Pipe-Friendly Framework for Basic Statistical Tests. R package version 0.7.2

Kaur-Bhambra, J., Wardak, D. L. R., Prosser, J. I., & Gubry-Rangin, C. (2022). Revisiting plant biological nitrification inhibition efficiency using multiple archaeal and bacterial ammonia-oxidising cultures. Biology and Fertility of Soils, 58(3), 241–249. 10.1007/s00374-020-01533-1

Müller, K., Wickham, H. (2023). _tibble: Simple Data Frames_. R package version 3.2.1. https://CRAN.R-project.org/package=tibble

Oksanen, J., Simpson, G., Blanchet, F., Kindt, R., Legendre, P., Minchin, P., O’Hara, R., Solymos, P., Stevens, M., Szoecs, E., Wagner, H., Barbour, M., Bedward, M., Bolker, B., Borcard, D., Carvalho, G., Chirico, M., De Caceres, M., Durand, S., Evangelista, H., FitzJohn, R., Friendly, M., Furneaux, B., Hannigan, G., Hill, M., Lahti, L., McGlinn, D., Ouellette, M., Ribeiro Cunha, E., Smith, T., Stier, A., Ter Braak, C., Weedon, J. (2022). _vegan: Community Ecology Package_. R package version 2.6–4, https://CRAN.R-project.org/package=vegan

Papadopoulou, E. S., Bachtsevani, E., Lampronikou, E., Adamou, E., Katsaouni, A., Vasileiadis, S., … Karpouzas, D. G. (2020). Comparison of Novel and Established Nitrification Inhibitors Relevant to Agriculture on Soil Ammonia- and Nitrite-Oxidizing Isolates. Frontiers in Microbiology, 11. 10.3389/fmicb.2020.581283

Pedersen, T. (2024). patchwork: The Composer of Plots. R package version 1.2.0

Powell, S. J., & Prosser, J. I. (1986). Effect of copper on inhibition by nitrapyrin of growth of Nitrosomonas europaea. Current Microbiology, 14(3), 177–179. 10.1007/bf01568371

R Core Team (2020). R: A Language and Environment for Statistical Computing. Austria: R Foundation for Statistical Computing

Ritz, C., Baty, F., Streibig, J. C., and Gerhard, D. (2016). Dose-response analysis using R. PLoS One 10:e0146021. 10.1371/journal.pone.0146021

Shaw, L. J., Nicol, G. W., Smith, Z., Fear, J., Prosser, J. I., & Baggs, E. M. (2006). Nitrosospira spp. can produce nitrous oxide via a nitrifier denitrification pathway. Environmental Microbiology, 8(2), 214–222. 10.1111/j.1462-2920.2005.00882.x

Wickham, H. ggplot2: Elegant Graphics for Data Analysis. Springer-Verlag New York, 2016.

Wickham, H., Bryan, J. (2023). _readxl: Read Excel Files_. R package version 1.4.3. https://CRAN.R-project.org/package=readxl

Wickham, H., François, R., Henry, L., Müller, K., Vaughan, D. (2023). _dplyr: A Grammar of Data Manipulation_. R package version 1.1.4. https://CRAN.R-project.org/package=dplyr

